# PEP444c encoded within the *MIR444c* gene regulates microRNA444c accumulation in barley

**DOI:** 10.1101/2023.05.24.542045

**Authors:** A Chojnacka, A Smoczynska, D Bielewicz, A Pacak, G Hensel, J Kumlehn, WM Karlowski, M Grabsztunowicz, E Sobieszczuk-Nowicka, A Jarmolowski, Z Szweykowska-Kulinska

## Abstract

MicroRNAs are small, non-coding RNA molecules that regulate expression of their target genes. The *MIR444* gene family is present exclusively in monocotyledons, and microRNAs444 from this family have been shown to target certain MADS-box transcription factors in rice and barley. We identified three barley *MIR444* (*MIR444a*/*b*/*c*) genes and comprehensively characterized their structure and the processing pattern of the primary transcripts (pri-miRNAs444). Pri-microRNAs444 undergo extensive alternative splicing, by which functional and non-functional pri-miRNA444 isoforms are generated. We show that barley pri-miRNAs444 contain numerous open reading frames (ORFs) whose transcripts associate with ribosomes. Using specific antibodies, we provide evidence that selected ORFs encoding PEP444a within *MIR444a* and PEP444c within *MIR444c* are expressed in barley plants. Moreover, we demonstrate that CRISPR-associated endonuclease 9 (Cas9)-mediated mutagenesis of the PEP444c encoding sequence results in a decreased level of *PEP444* transcript in barley shoots and roots, and a 5-fold reduced level of mature microRNA444c in roots. Taken together, our observations suggest that PEP444c encoded by the *MIR444c* gene is involved in microRNA444c biogenesis in barley.

## INTRODUCTION

MicroRNAs are small, single-stranded RNA molecules that regulate expression of their target genes which themselves are involved in developmental processes or response to environmental stimuli (Sunkar et al., 2012; Bartel, 2009; Voinnet, 2009). Plant *MIR* genes are transcribed by RNA Pol II and primary transcripts (pri-miRNAs) form a characteristic stem-loop structure containing a microRNA/microRNA* duplex (Reinhart et al., 2002; Rogers and Chen, 2013). Pri-miRNAs undergo maturation processes that involve splicing, alternative splicing, nucleotide modifications and alternative polyadenylation site selection (Bielewicz et al., 2012; Yan et al., 2012; Jia and Rock, 2013). In the nucleus, microRNA and microRNA* are excised from the stem-loop structure in a two-step process by RNase III enzyme DICER-LIKE (DCL1), which is a key component of microprocessor machinery (Vazquez et al., 2004; Li et al., 2005; Yu et al., 2005; Kurihara et al., 2006; Lobbes et al., 2006; Laubinger et al., 2008; Yang et al., 2010; Manavella et al., 2012; Ren et al., 2012; Zhan et al., 2012). In the final stage, the microRNA is loaded into the AGO1 protein in the nucleus and exported to the cytoplasm to execute posttranscriptional gene regulation by cleaving the target mRNA or inhibiting translation (Vaucheret et al. 2004; Baumberger and Baulcombe 2005; Bologna et al. 2018).

One of the exciting phenomena concerning microRNA biogenesis regulation is the discovery of short open reading frames (ORFs) encoding functional peptides (miPEPs) within *MIR* genes (Lauressergues et al., 2015). These miPEPs are encoded in the 5’ region of numerous pri-miRNAs, with this coding sequence always being located upstream of the microRNA’s stem-loop structure. miPEPs were shown to stimulate biogenesis of their cognate microRNAs, which results in a substantial increase of microRNA level and, consequently, downregulation of target gene expression (Lauressergues et al., 2015; Couzigou et al., 2016; Chen et al., 2020, Sharma et al., 2020; Zhang et al., 2020). Until now, miPEPs were discovered in *Arabidopsis thaliana*, *A. lyrata, Medicago truncatula*, *Glycine max*, *Brassica rapa,* and *Vitis vinifera* (Lauressergues et al., 2015; Combier et al., 2020; Couzigou and Combier, 2016; Sharma et al., 2020; Sharma et al., 2016; Nag et al., 2009; Cheng et al., 2016; Morozov et al., 2019). It has been demonstrated that using artificial miPEP165a in *A. thaliana* positively affects the development of roots (Carlsbecker et al., 2010; Lauressergues et al., 2015). miPEP172c and miPEP167c have been shown to stimulate the formation of root nodules in legumes (Couzigou and Combier, 2016; Combier et al., 2018). Similarly, miPEP171d1 stimulates the formation of adventitious roots in *V. vinifera.* All these data show that miPEPs may be powerful tools used in propagating economically important plants (Chen et al., 2020). Until now, miPEPs were reported to be present in dicot plants. To our knowledge, there is thus far only one case where the presence of a miPEP was reported in a representative of the monocots, namely maize (Combier et al., 2016).

Members of the *MIR444* gene family have been found exclusively in monocot species such as *Oryza sativa, Triticum aestivum*, *Hordeum vulgare*, *Zea mays*, *Sorghum bicolor*, *Brachypodium* species, and *Saccharum officinarum* (Sunkar et al., 2005; Sunkar et al., 2008). The characteristic feature of *MIR444* genes is the presence of an intron that separates two halves of the microRNA precursor structure. One half containing microRNA* is situated in the upstream exon while the other one carrying microRNA, resides in the proximal downstream exon. Consequently, intron removal is an essential step in forming the stem-loop structure of pri-miRNA444 from which microRNA444 can be produced. It has been reported that some MADS-box family transcription factors are encoded by the opposite strand of MIR444 genes and consequently contain target sequences for cognate microRNAs444 (Sunkar et al., 2005; Lu et al., 2008). In rice, microRNAs of the *MIR444* family regulate the expression of four MADS-box transcription factors. These are encoded by *OsMADS23*, *OsMADS27a*, *OsMADS27b,* and *OsMADS57* and involved in the plants’ responses to NO_3_ ^-^, P and NH ^+^ supply and to the rice stripe virus (RSV) (Jiao et al., 2020; Wang et al., 2016; Yan et al., 2014). Furthermore, microRNA444a, by regulating the level of its cognate transcription factor OsMADS57, controls the level of the DWARF14 protein, which is a negative regulator of tillering in rice (Guo et al., 2013). Therefore, microRNAs belonging to the *MIR444* family play essential roles in growth and development of rice. However, our knowledge about *MIR444* genes and their associated microRNAs is very limited in other monocot species. Barley (*H. vulgare*) is a crop plant of great economic importance. Almost no information exists on barley *MIR444* gene number, microRNA444 activity, and their targets. In the present investigation, we identify three barley *MIR444s*. We show complex pri-miRNA444s processing and the identification of potential ORFs. Some of them may encode miPEPs. Moreover, we reveal that pri-miRNAs from the *MIR444* family are actively associated with ribosomes. In the case of the *MIR444c* gene, we show that the formation of truncated miPEP (PEP444c) encoded by this gene entails a decreased level of microRNA444c. To our knowledge, this is the first evidence of miPEP existence and its impact on microRNA accumulation in monocot plants.

## RESULTS

### Identification of cis-regulatory elements in promoter regions of barley *MIR444* **genes**

Bioinformatic analysis of the barley genome revealed the presence of three *MIR444* genes: *MIR444a*, *MIR444b,* and *MIR444c*. To learn more about these genes’ biological function, we used The New PLACE database to analyze cis-regulatory elements (CREs) in the promoter sequences (Higo et al., 1999). We found multiple CREs involved in abiotic stress response (Table S1). The most abundant CRE in all three analyzed *MIR444* genes is CACTFTPPCA1, a motif associated with the regulation of nitrogen metabolism in plants (Bai et al., 2013). Among the others, the most numerous regulatory motifs are those involved in the plant response to light (Tiwari et al., 2016; Hudson and Quail, 2003; Villain et al., 1996; Terzaghi and Cashmore, 1995). In addition, we identified CREs associated with the response to heat stress and copper (Belity et al., 2022; Tiwari et al., 2016; Rieping and Schöffl, 1992).

Considering the abundance of the regulatory elements involved in the regulation of nitrogen metabolism that we found in promoter regions of all barley *MIR444* genes, we analyzed the level of pri-miRNA444a/b/c in 2-week-old barley plants under control and nitrogen excess stress conditions. We amplified pri-miRNA444a/b/c fragments encompassing the stem-loop structure region. Whereas the level of pri-miRNA444a proved not to be affected by nitrogen excess, the level of pri-miRNA444b was significantly increased in barley roots under the stress condition, with its level not being affected in shoots. The level of pri-miRNA444c in roots was neither affected by nitrogen excess stress. Of note, pri-miRNA444c was not detectable whatsoever in barley shoots under both analyzed conditions (Figure 1A, 1B). sRNA seq analysis performed in barley roots and shoots also did not reveal significant changes in the levels of mature microRNA444a/b/c (Figure 1C, 1D). The expression of the *NRT1.1* (*NITRATE TRANSPORTER 1*) was used as a marker of nitrogen excess stress (Smoczynska et al., 2022).

**Figure 1.**
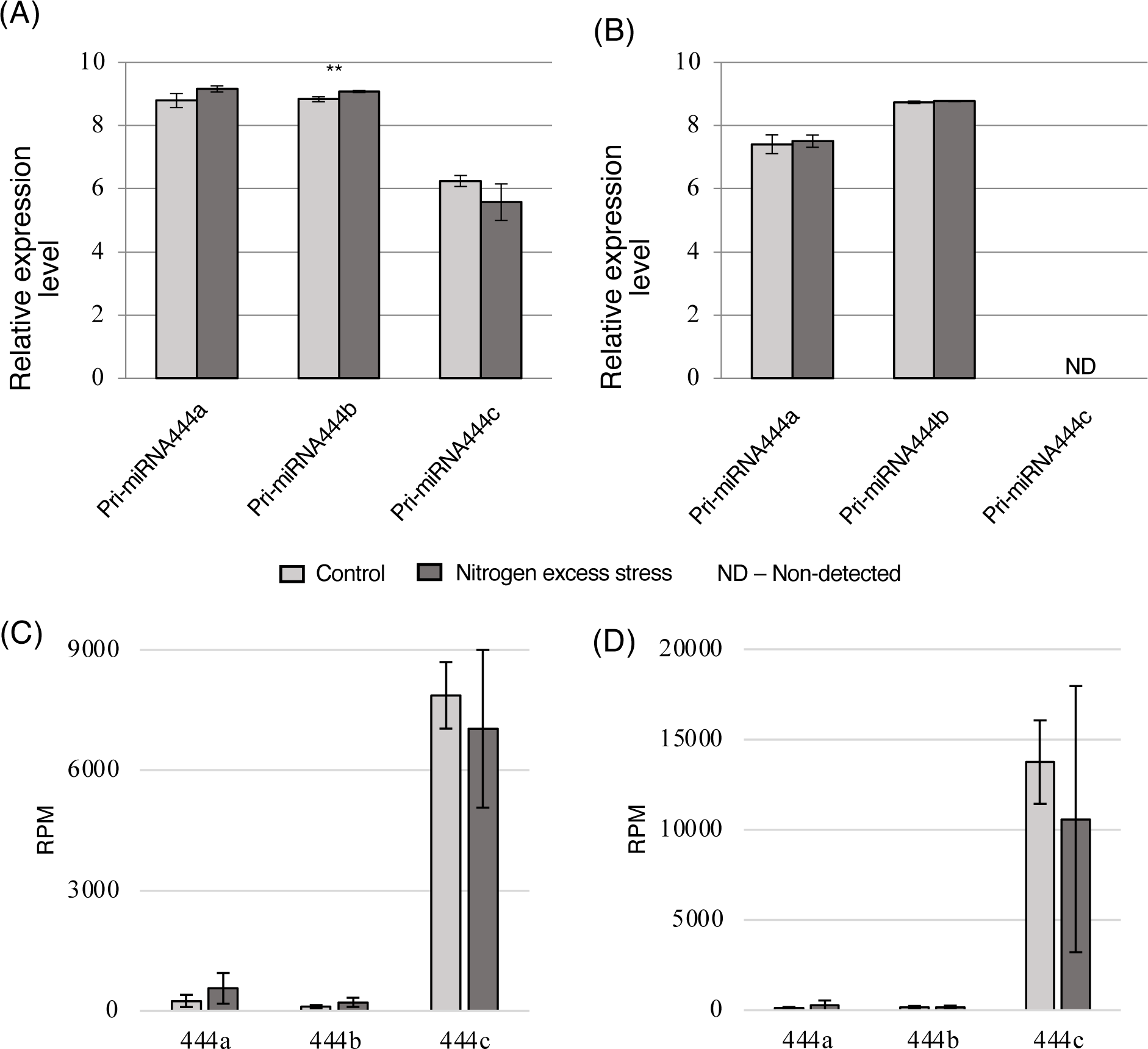
Relative expression levels of pri-miRNA444a/b/c assessed in (A) roots and (B) shoots, and mature microRNA444a/b/c in (C) roots and (D) shoots under control and nitrogen excess stress performed with the use of RT-qPCR and sRNA seq. Light grey bars represent control conditions, and dark grey bars represent nitrogen excess stress conditions. (A, B) Values are shown as the mean ± SD (n=3) from three independent experiments; ND – not detectable. Error bars (n=3) indicate SD, and asterisks indicate a significant difference between the sample and control (\*\**P* < 0.01). (C, D). The number of normalized counts for three biological replicates (RPM, reads per million) is shown.

### *MIR444* genes in barley are characterized by complex exon-intron structure

To determine the structure of *MIR444a*/b/*c* genes and their transcripts, we performed 5’ and 3’ RACE analyses. The length of the pri-miRNA was calculated based on the longest pri-miRNA 5’ and 3’ RACE products. To reveal that the longest pri-miRNA 5’ and 3’ ends belong to the same precursor, we conducted RT-PCR for all three analyzed pri-miRNAs using primers designed for the 5’ and 3’ ends of the most prolonged RACE products. The intron-exon structure of barley *MIR444* genes was established by alignment of the pri-miRNA sequences with genomic sequences deposited in the Ensembl Plants and NCBI databases. Our analysis revealed a complex exon-intron architecture of all three *MIR444* genes in barley. The lengths of individual exons and introns identified in the *MIR444a/b/c* genes are presented in Table 1.

**Table 1.** The length and structure of three characterized *MIR444* genes in *H. vulgare*.

| *MIR* gene | Chromosome | Position (strand) | Length (bp) | Number of exons (length) | Number of introns (length) |
| --- | --- | --- | --- | --- | --- |
| *444a* | 6 | 490703243-490708100 (-) | 4858 | 6 (96, 96, 104, 113, 70, 520) | 5 (1142, 1940, 572, 71, 134) |
| *444b* | 7 | 117611827-117622532 (+) | 10706 | 5 (240, 62, 109, 66, 445) | 4 (1019, 2378, 130, 6257) |
| *444c* | 2 | 512343713-512352484 (-) | 8772 | 5 (569, 57, 97, 74, 454) | 4 (3245, 3204, 992, 80) |

The *MIR444a* gene (genomic position 6H:490703243-490708100) comprises six exons and five introns. Using full transcript analysis, we found that pri-miRNA444a undergoes extensive alternative splicing events generating isoforms that can be divided into two groups. Accordingly, five functional (pri-miRNA444a.1-5) and three non-functional (pri-miRNA444a.6-8) isoforms were distinguished (Figure 2). Moreover, 3’ RACE analysis revealed the presence of six alternative poly(A) sites located in these miRNAs’ last exons. The expression levels of four out of eight identified isoforms (two functional: pri-miRNA444a.1 and pri-miRNA44a.5, and two non-functional isoforms: pri-miRNA444a.7 and pri-miRNA444a.8) was analysed in detail under control and excess nitrogen conditions. There was no significant difference between the expression levels of functional isoforms under control and nitrogen excess stress conditions. However, the non-functional isoform pri-miRNA444a.7 was detectable only in roots upon nitrogen excess stress. Another non-functional isoform (pri-miRNA444a.8) was detectable in roots, but its level was not affected by the conditions tested. Whereas it was also detected in shoots under control conditions, its level was dramatically decreased (i.e. not detectable) upon excess nitrogen (Figure 3). Taken together, our analyses reveal dynamic fluctuations in the presence of pri-miRNA444a non-functional isoforms in shoots and roots in dependence of exposure to excess nitrogen stress.

**Figure 2.**
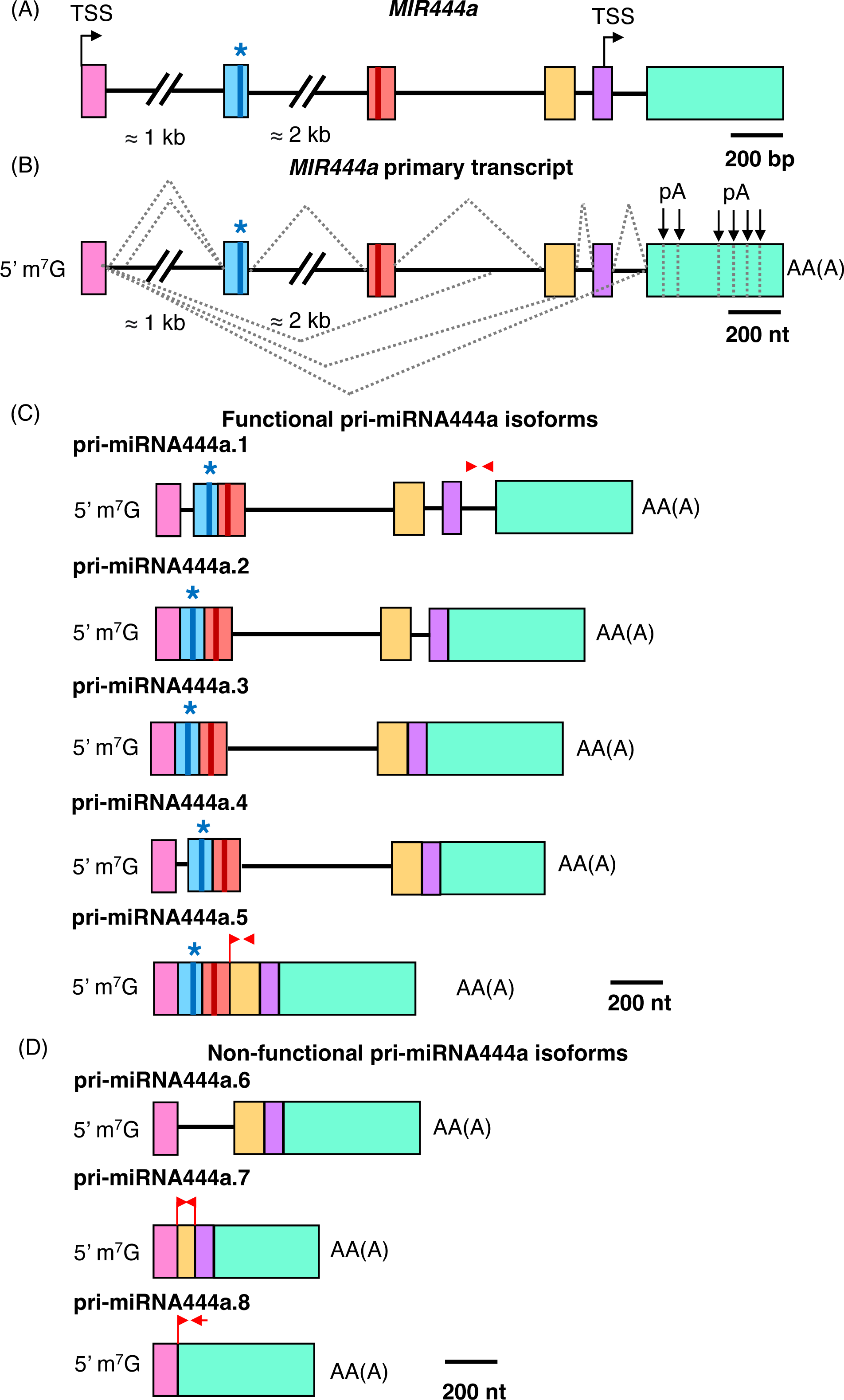
*MIR444a* gene structure, its primary transcript, and processed pri-miRNA444a isoforms. (A) *MIR444a* gene structure; arrows indicate identified transcription start sites (TSS). The scale bar represents 200 base pairs (bp). (B) Primary transcript structure of *MIR444a* gene. Gray dotted lines represent identified splicing events. (C) Functional pri-miRNA444a isoforms; red arrows indicate primer positions used in RT-qPCR experiments. (D) Non-functional pri-miRNA444a isoforms; red arrows indicate primer positions used in RT-qPCR experiments. (A-D) Colored boxes and black lines correspond to exons and introns, respectively. (B-D) m^7^G -7-methylguanosine; arrows tagged as pA depict identified polyadenylation sites; AA(A) depicts polyA tail. The scale bar represents 200 nucleotides (nt). (A-C) Blue bar and blue star represent microRNA* position, and red bar indicates microRNA position. (C-D) pri-miRNA444a.1-pri-miRNA444a.8 represent detected pri-miRNA444a isoforms.

**Figure 3.**
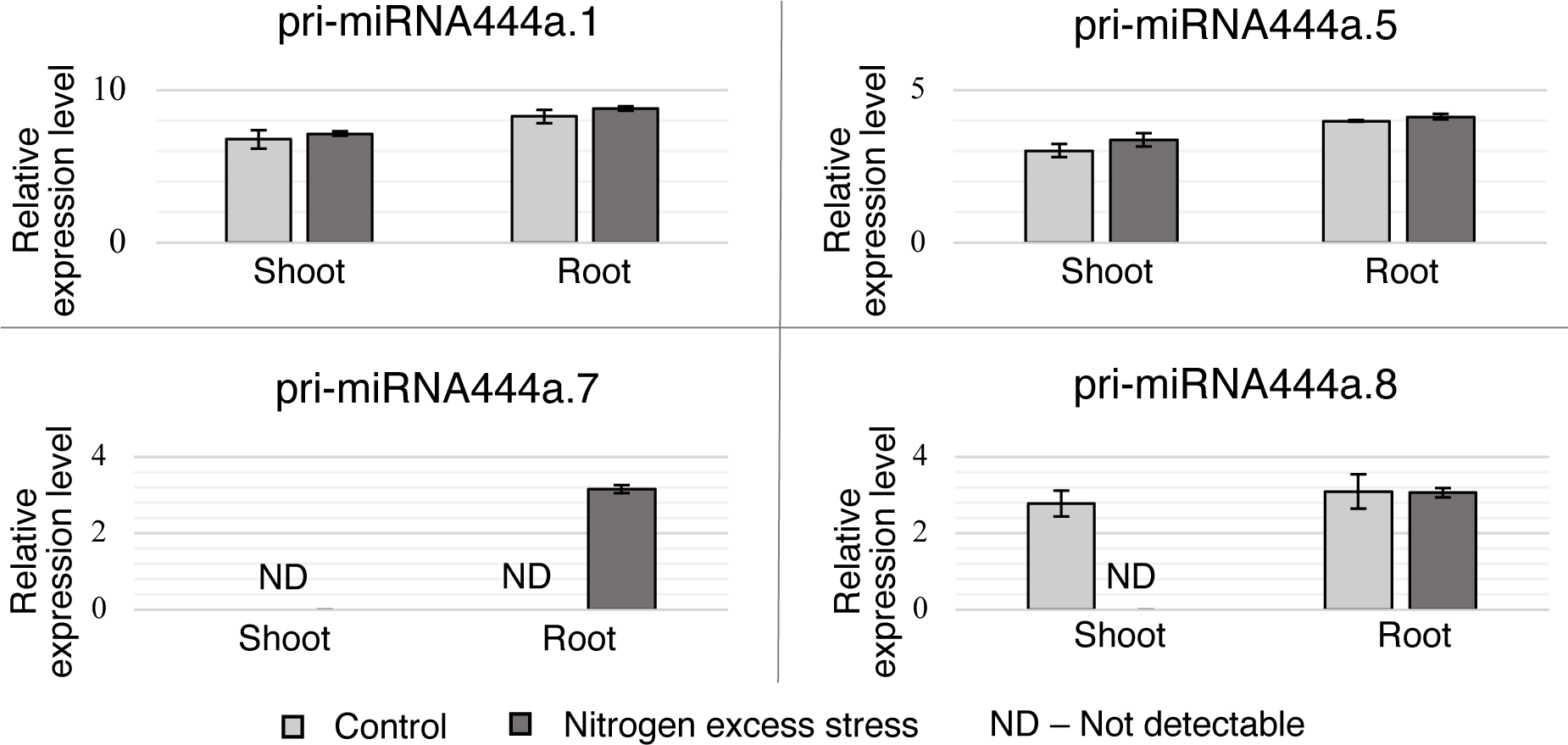
Pri-miRNA444a of the non-functional but not of the functional isoforms show dynamic fluctuations in shoots and roots under nitrogen excess stress. RT-qPCR measurements of detectable pri-miRNA444a isoforms under control and nitrogen excess stress. Light grey bars represent control conditions, and dark grey bars show nitrogen excess stress conditions; error bars indicate SD (n=3). ND – not detectable.

5’ RACE analyses revealed an alternative transcription start site (TSS) located in the fifth exon of the *MIR444a* gene. However, *in silico* analysis of the 1 kb region upstream of this non-canonical TSS detected the presence of numerous promoter elements but no classical basic promoter motifs like TATA box or initiator element.

The *MIR444b* gene (genomic localization: 7H:117611827-117622532) consists of five exons and four introns. Like pri-miRNA444a, also pri-miRNA444b undergoes alternative splicing and generates multiple isoforms. We detected four functional (pri-miRNA444b.1-4) and one non-functional isoform (pri-miRNA444b.5) (Figure 4). We also established the presence of five alternative poly(A) sites in the last exon. We performed quantitative expression analyses of three out of the five pri-miRNA444b isoforms, namely the two functional pri-miRNAs 444b.1 and 2 and the non-functional pri-miRNA444b.5 under control and nitrogen excess conditions. The level of pri-miRNA444b.1 was slightly increased in shoots under nitrogen excess, while it was unchanged in roots under both analyzed conditions. The second studied functional isoform was unchanged in both shoots and roots under control and nitrogen excess. The non-functional isoform was slightly decreased in roots under nitrogen excess, whereas it was unaltered in shoots under both analyzed conditions (Figure 5).

**Figure 4.**
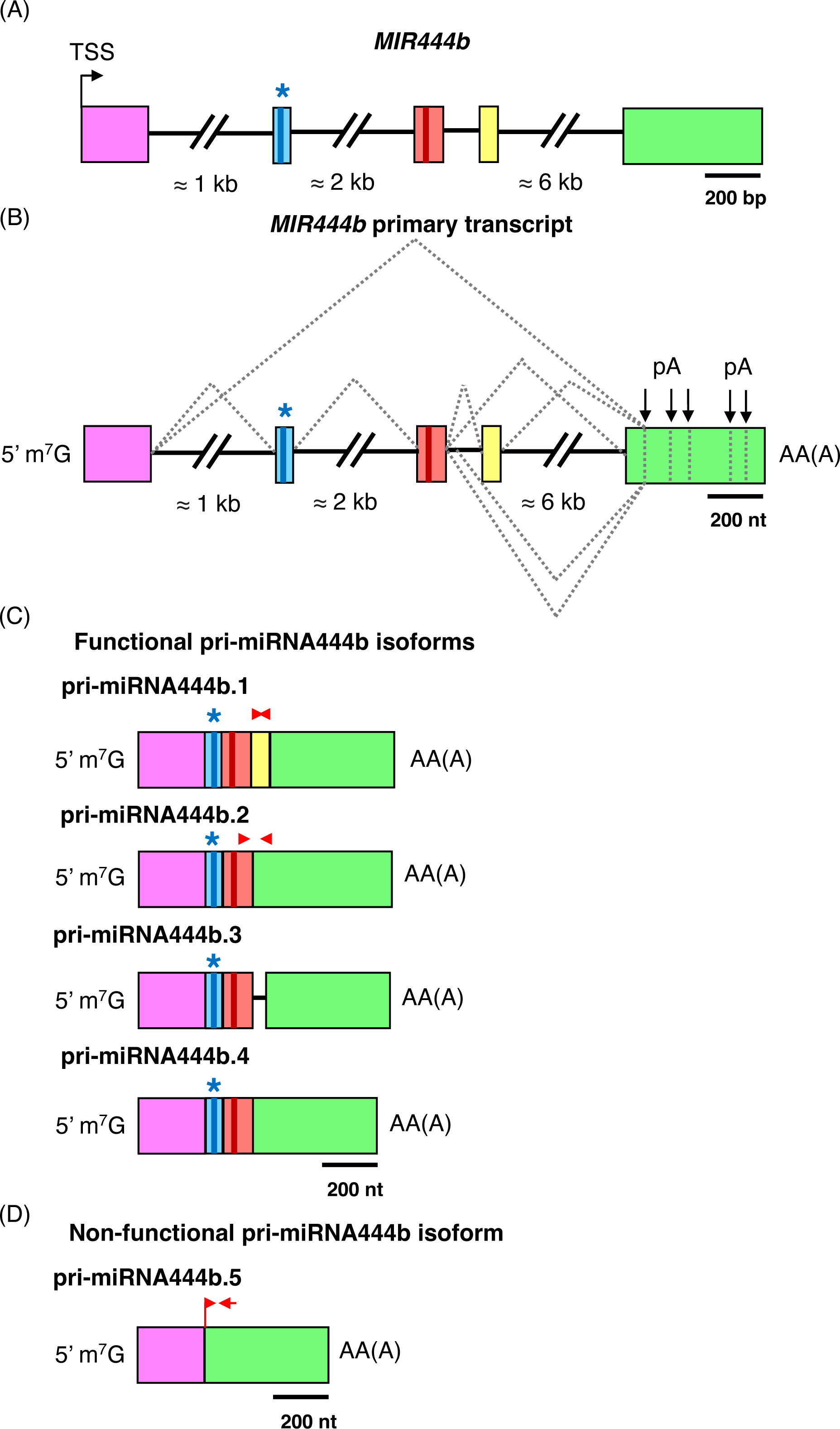
*MIR444b* gene structure, its primary transcript, and processed pri-miRNA444b isoforms. (A) *MIR444b* gene structure; (B) pri-miRNA444b structure; (C) functional pri-miRNA444b; (D) non-functional pri-miRNA444b isoforms. Details are marked in the same way as in Figure 2. (C-D) pri-miRNA444b.1-pri-miRNA444b.5 represent detected pri-miRNA444b isoforms.

**Figure 5.**
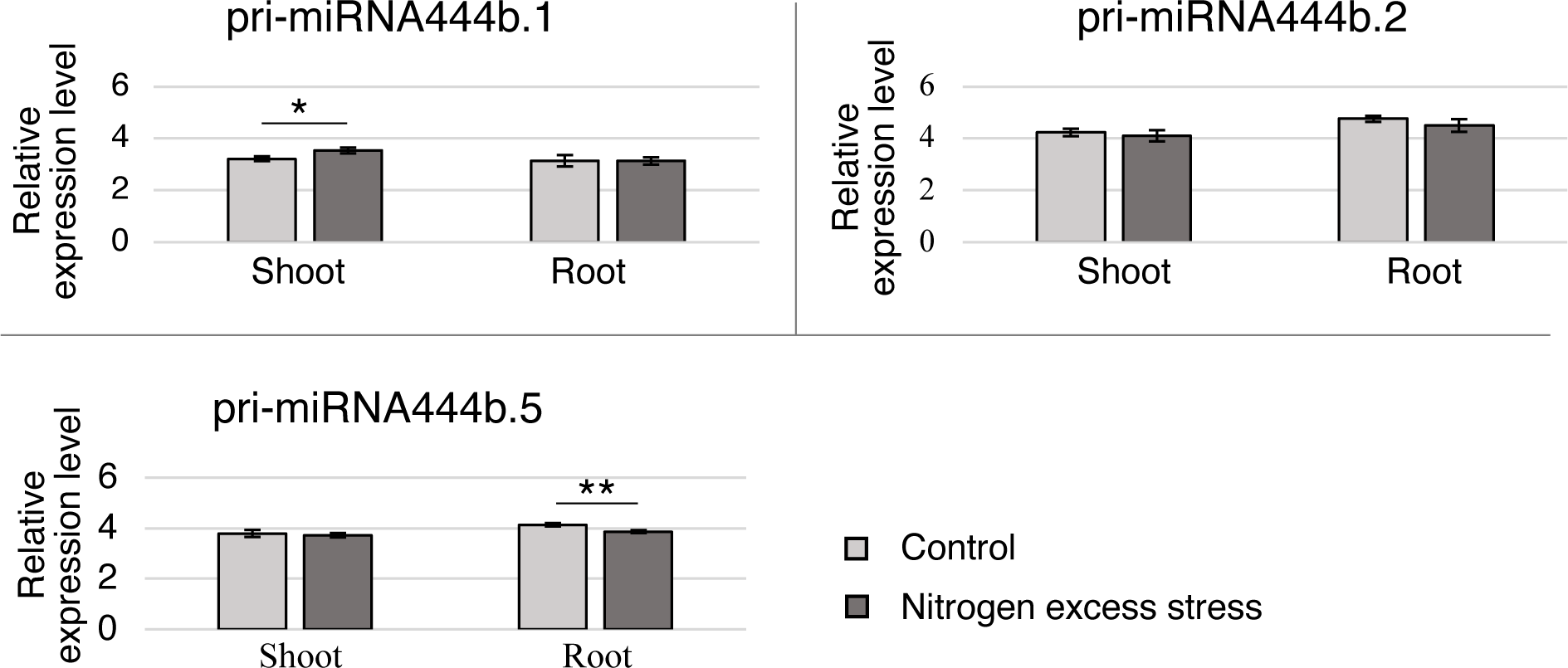
The level of detectable pri-miRNA444b isoforms is not affected by nitrogen stress. RT-qPCR measurements of selected pri-miRNA444b isoforms under control and nitrogen excess stress. Light grey bar represents control conditions, and dark grey bar shows nitrogen excess stress conditions. Error bars indicate SD (n=3); asterisks indicate a significant difference between the sample and control (*P < 0.05; **P < 0.01). The level of detectable pri-miRNA444b isoforms is not affected by nitrogen stress.

The *MIR444c* gene (genomic position 2H:512343713-512352484) contains five exons and four introns. In the first exon, we identified two alternative TSS residing in closed vicinity. Pri-miRNA444c, similarly to pri-miRNA444a and pri-miRNA444b, undergoes alternative splicing. We detected one functional and four non-functional isoforms, specifically pri-miRNA444c.1 and pri-miRNA444c.2-5, respectively (Figure 6). Interestingly, 3’ RACE analyses revealed the presence of five alternative poly(A) sites in the first intron (three poly(A) sites), second intron (one poly(A) site), third intron (one poly(A) site) as well as in the last exon (two poly(A) sites). We analyzed the expression profile of the functional (pri-miRNA444c.1) and two non-functional isoforms (pri-miRNA444c.2 and pri-miRNA444c.4) under control and nitrogen excess stress conditions. Pri-miRNA444c.1 was detected only in roots at the same level under both conditions studied, while the other two isoforms showed no differences in their expression levels in shoots and roots (Figure 7).

**Figure 6.**
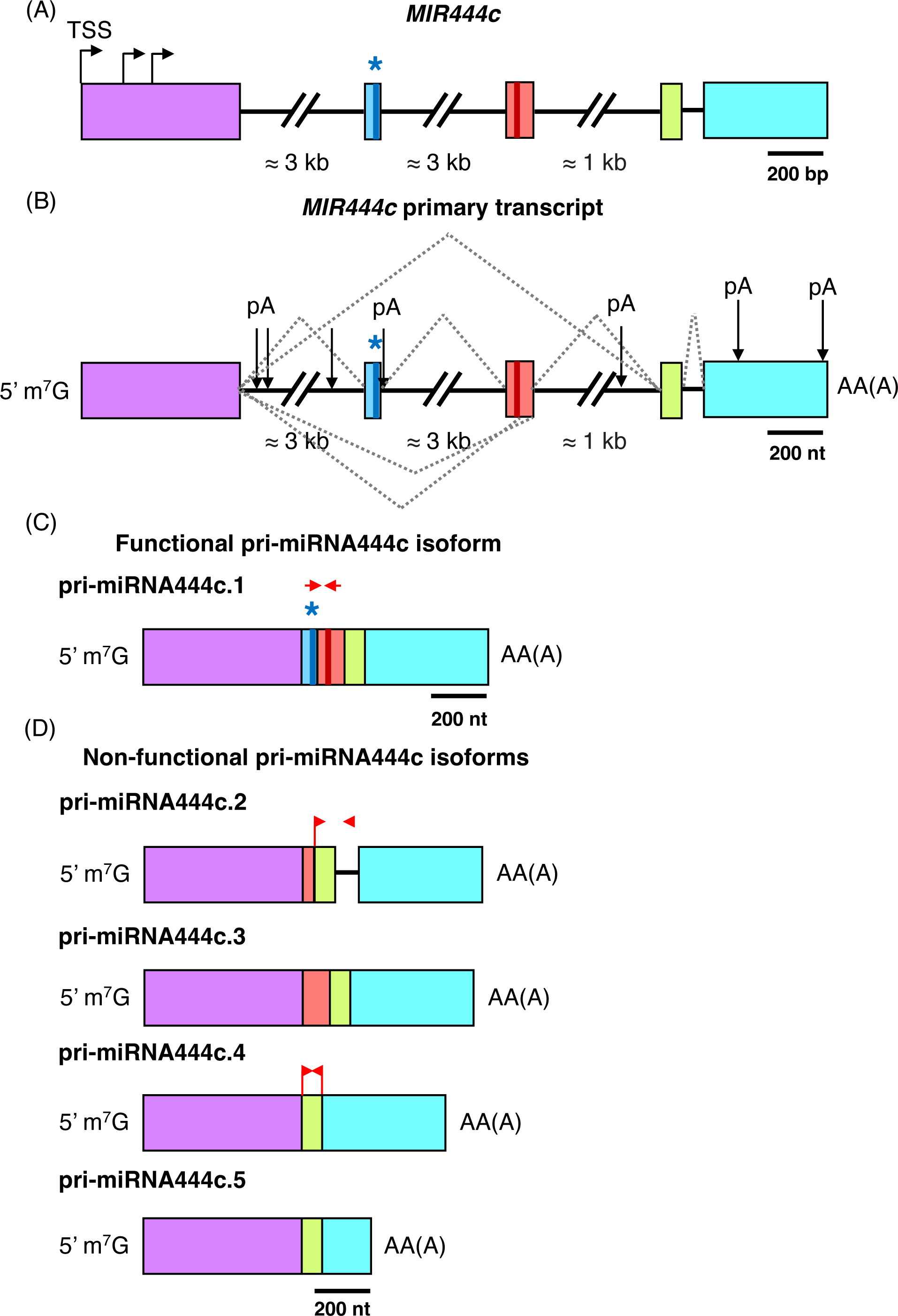
*MIR444c* gene structure, its primary transcript, and processed pri-miRNA444c isoforms. (A) *MIR444c* gene structure; (B) pri-miRNA444c structure; (C) functional pri-miRNA444c; (D) non-functional pri-miRNA444c isoforms. Details in the Figure are marked in the same way as in Figure 2. (C-D) pri-miRNA444c.1-pri-miRNA444c.5 represent detected pri-miRNA444c isoforms.

**Figure 7.**
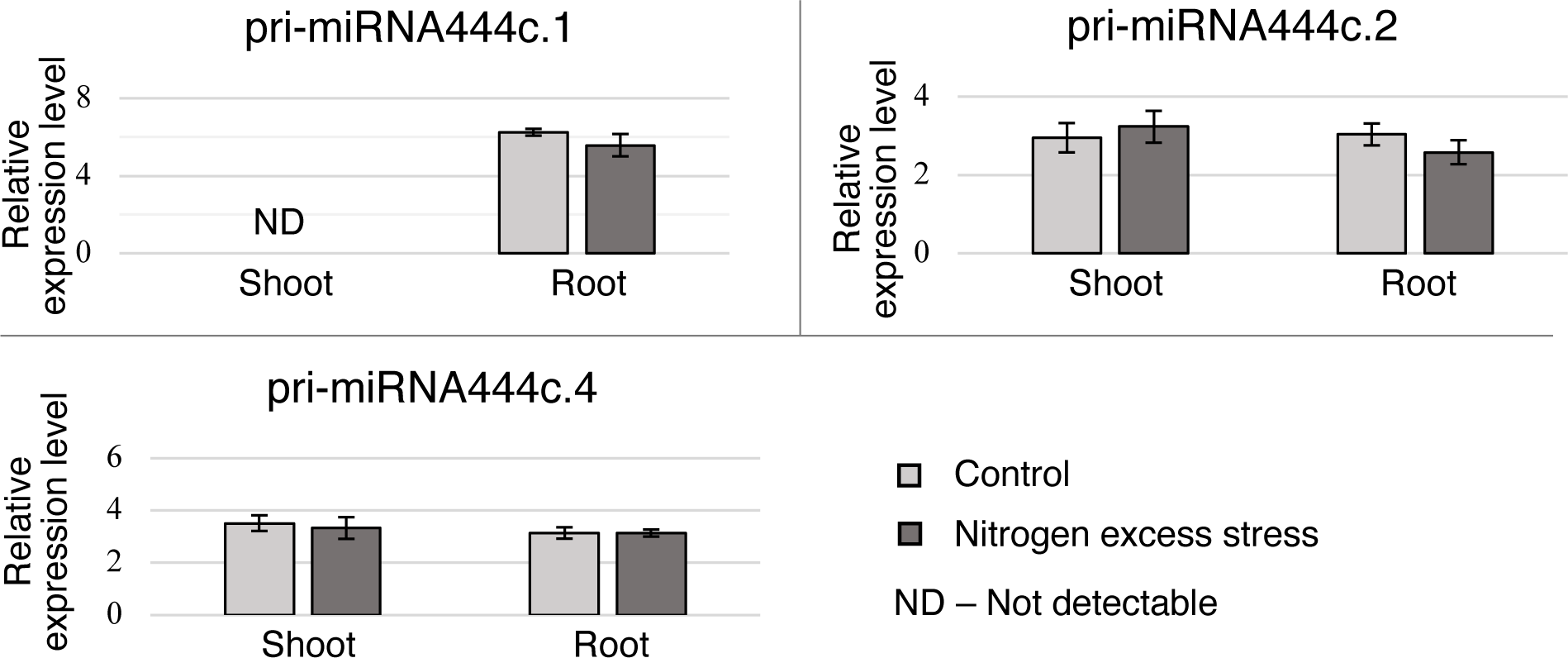
The level of the functional but not of the non-functional pri-miRNA444c isoforms is affected by nitrogen excess stress. RT-qPCR measurements of detectable pri-miRNA444c isoforms under control and nitrogen excess stress. Light grey bars represent control conditions, and dark grey bars show nitrogen excess stress conditions. Error bars indicate SD (n=3); ND –not detectable.

Summing up, our results concerning the representation of various pri-miRNA444a/b/c isoforms in organs and stress studied, we conclude that pri-miRNA444a and pri-miRNA444c show higher dynamics in pri-miRNA isoform levels than pri-miRNA444b.

### Pri-miRNAs from the *MIR444* gene family contain ORFs and associate with ribosomes

Using ORF Finder (https://www.ncbi.nlm.nih.gov/orffinder/) pri-miRNA444 sequences were analysed for the presence of ORFs (Stothard, 2000). Considering all pri-miRNA444a and pri-miRNA444b isoforms, six different ORFs were identified (1-6), and three in pri-miRNA444c, respectively (Table S2). For further analysis, an ORF encoding a peptide of 119 amino acids (aa) (PEP444a) located within the *MIR444a* gene and having its own TSS was selected. Another ORF encoding 51 aa (miPEP444b) located within the *MIR444b* gene and fulfilling all features allowing it to be defined as miPEP (short peptide, encoded upstream of the miRNA region bearing a stem-loop structure), and a third ORF encoding an 168 aa peptide located within the *MIR444c* gene that we called PEP444c, because it does not show all characteristics of a miPEP were selected, respectively. More specifically, it is encoded upstream of the region bearing stem-loop structure, while its size is beyond that of miPEPs (Lauressergues et al., 2015). The genomic location of the sequences encoding selected peptides is shown in Figure 8. Since PEP444a may have its own TSS, it can be considered as an independent transcription unit. However, we also identified one isoform of pri-miRNA444a (pri-miRNA444a.1) that can potentially encode this peptide. miPEP444b is encoded only in functional isoforms of the pri-miRNA444b, while PEP444c is encoded in all identified pri-miRNA444c isoforms.

**Figure 8.**
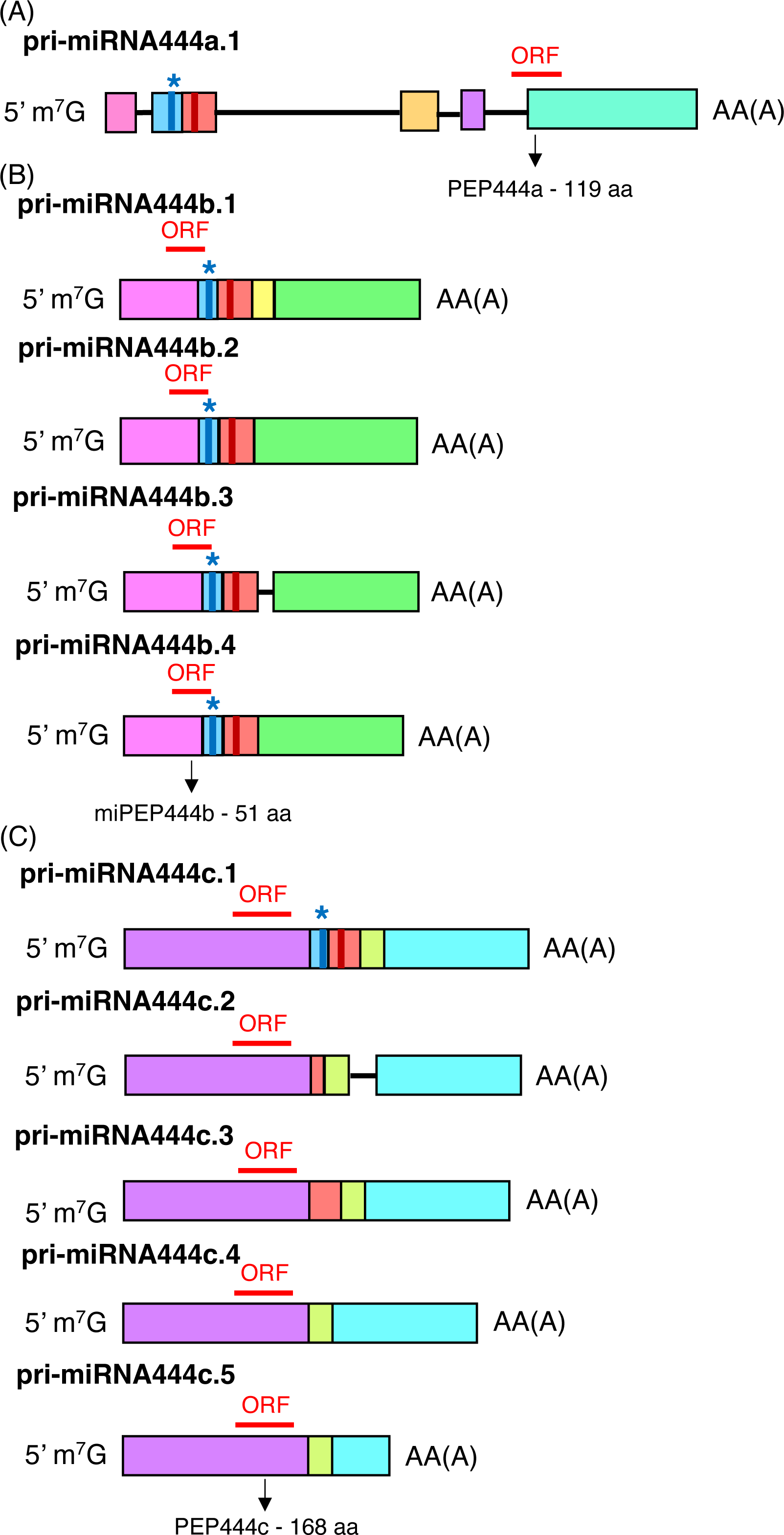
ORFs identified in pri-miRNAs444. Pri-miRNA444a (A), pri-miRNAs444b (B), and pri-miRNAs444c (C) with selected for further studies ORFs marked in red. The lengths of ORFs are given below each pri-miRNA444a/b/c; m^7^G -7-methylguanosine, pA - polyadenylation site.

To test whether the identified ORFs encoding putative peptides can be translated, polysome profiling using plant extracts isolated from 2-week-old barley shoots and roots grown under control and nitrogen excess stress conditions was performed. Cytoplasmic extracts were subjected to high-speed differential centrifugation in a sucrose density gradient (Mustroph et al., 2009). Supplementary Figure 1A shows an example of polysome gradient profiling. Supplementary Figure 1B and 1C show the association of *ARF1* (*ADP-ribosylation factor like-1,* GenBank: AJ508228.2) mRNA with ribosomes in roots and shoots, respectively. Specific RT-PCR products in polysomal fractions (marked with number 6-12) in both organs were observed. After EDTA treatment, which disrupts polysome association with mRNA, we detected the shift of RT-PCR products towards the fractions containing RNA not associated with ribosomes.

An analogous analysis was performed for the PEP444a coding sequence. We observed a specific association with ribosomes only in barley shoots under control and excess nitrogen conditions (Figure 9). The miPEP444b coding sequence analysis revealed the association with ribosomes mainly in barley roots under both tested conditions and in shoots of the control only (Figure 10). The sequence encoding PEP444c was associated with ribosomes in a specific manner in barley shoots and roots under control and nitrogen excess stress (Figure 11). Table 2 summarizes the association of the sequences encoding selected peptides with polysomes.

**Figure 9.**
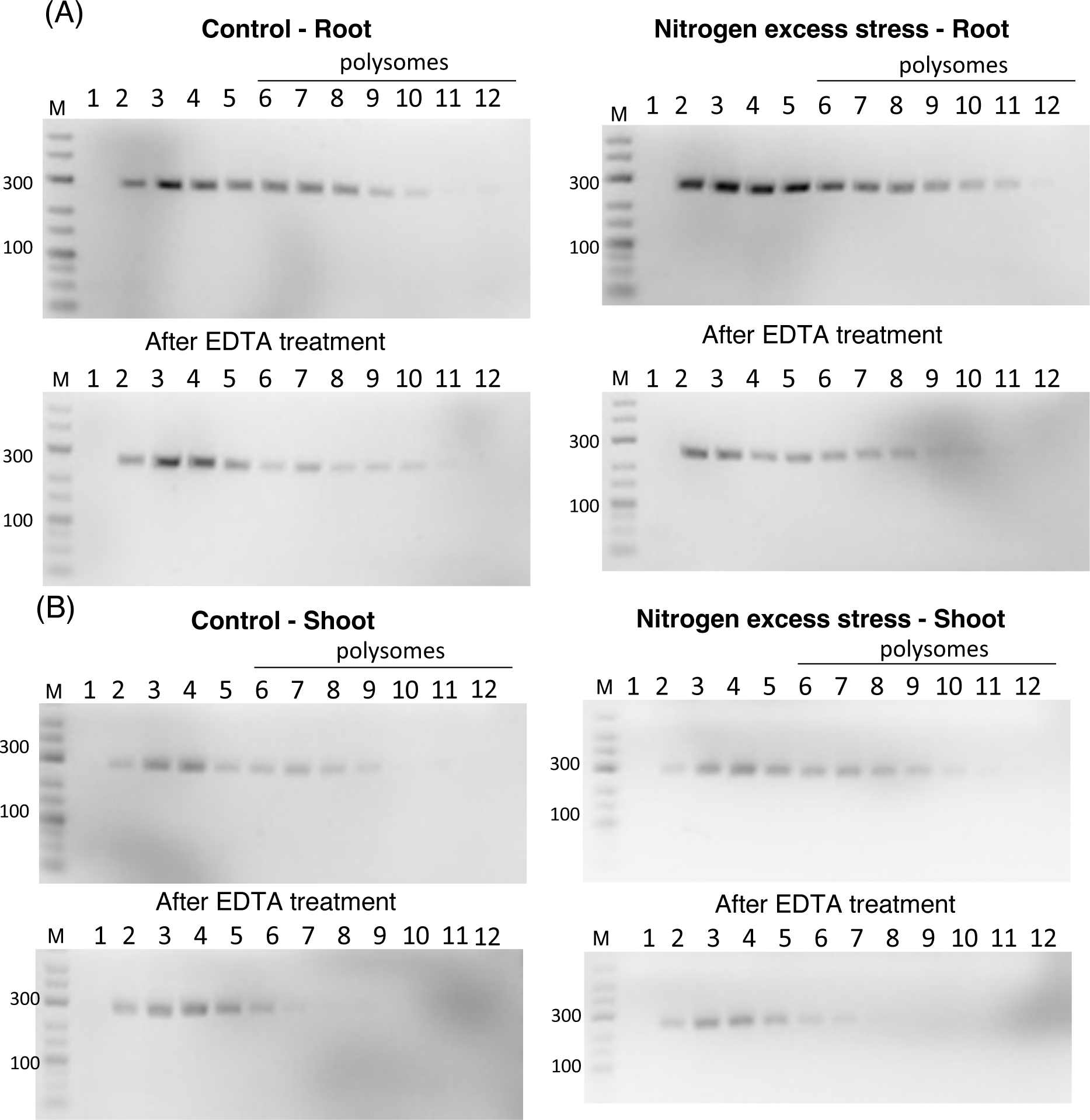
Pri-miRNA444a associates with ribosomes in a specific manner. RT-PCR analysis of the association of the pri-miRNA444a with ribosomes in barley roots (A) and shoots (B) under control and nitrogen excess stress. Numbers given above the agarose gels correspond to the numbers of the collected fractions. Polysomes as of the sixth fraction are marked. RT-PCR products observed of the sixth fraction indicate the specific binding of analysed pri-miRNA444a with ribosomes. After EDTA treatment, shift of RT-PCR products towards the fractions containing mRNA not associated to ribosomes is observed. Primers used for pri-miRNA444a amplification encompassed the region encoding PEP444a. M - GeneRuler Low-Range DNA Ladder.

**Figure 10.**
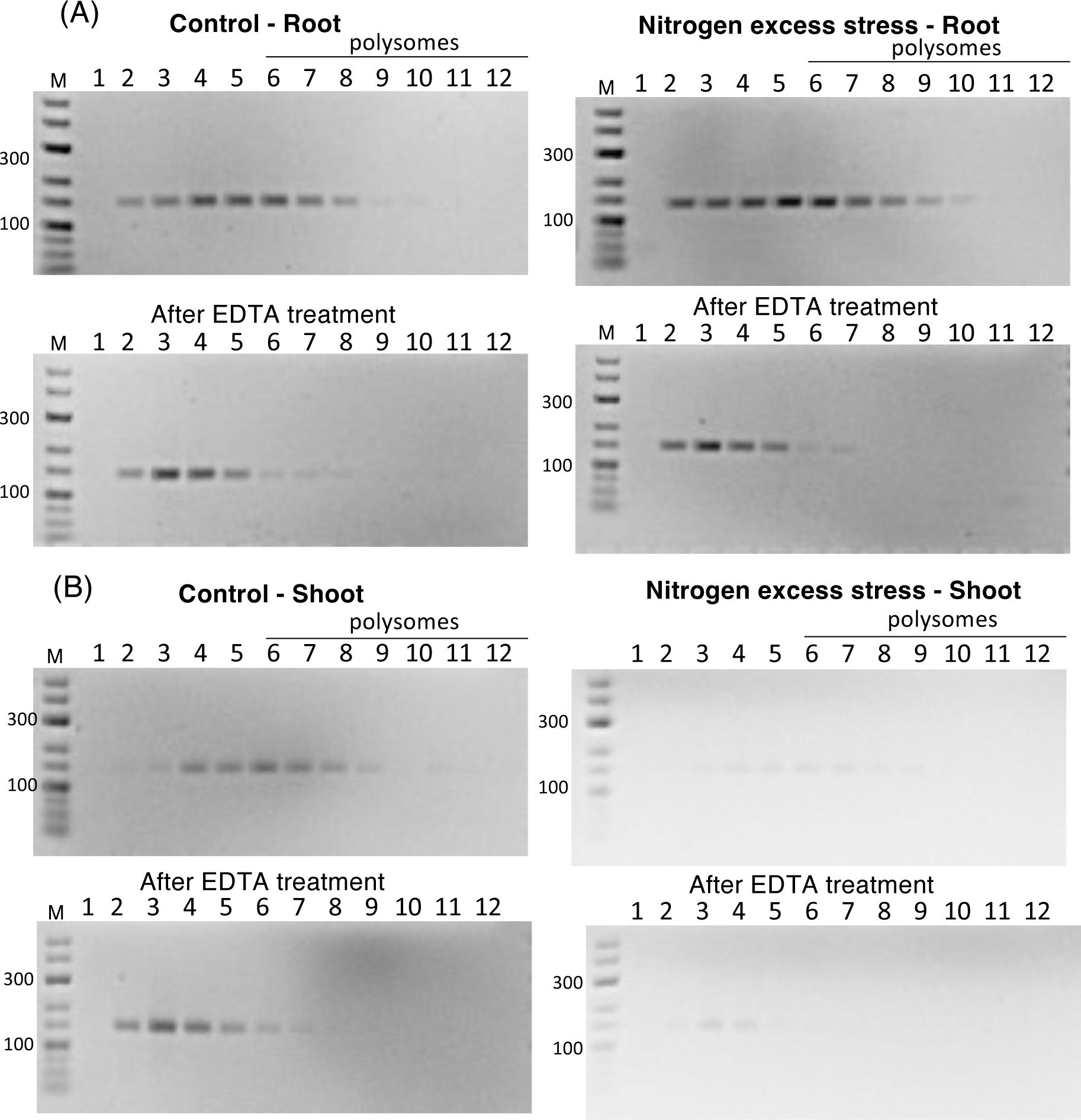
Pri-miRNA444b associates with ribosomes in a specific manner. RT-PCR analysis of the association of the pri-miRNA444b with ribosomes in barley roots (A) and shoots (B) under control and nitrogen excess stress conditions. Numbers above the agarose gels correspond to the numbers of the collected fractions. Polysomes as of the sixth fraction are marked. RT-PCR products observed of the sixth fraction indicate the specific binding of analysed pri-miRNA444b with ribosomes. After EDTA treatment, shift of RT-PCR products towards the fractions containing mRNA not associated to ribosomes is observed. Primers used for pri-miRNA444b amplification encompassed the region encoding miPEP444b. M - GeneRuler Low-Range DNA Ladder.

**Figure 11.**
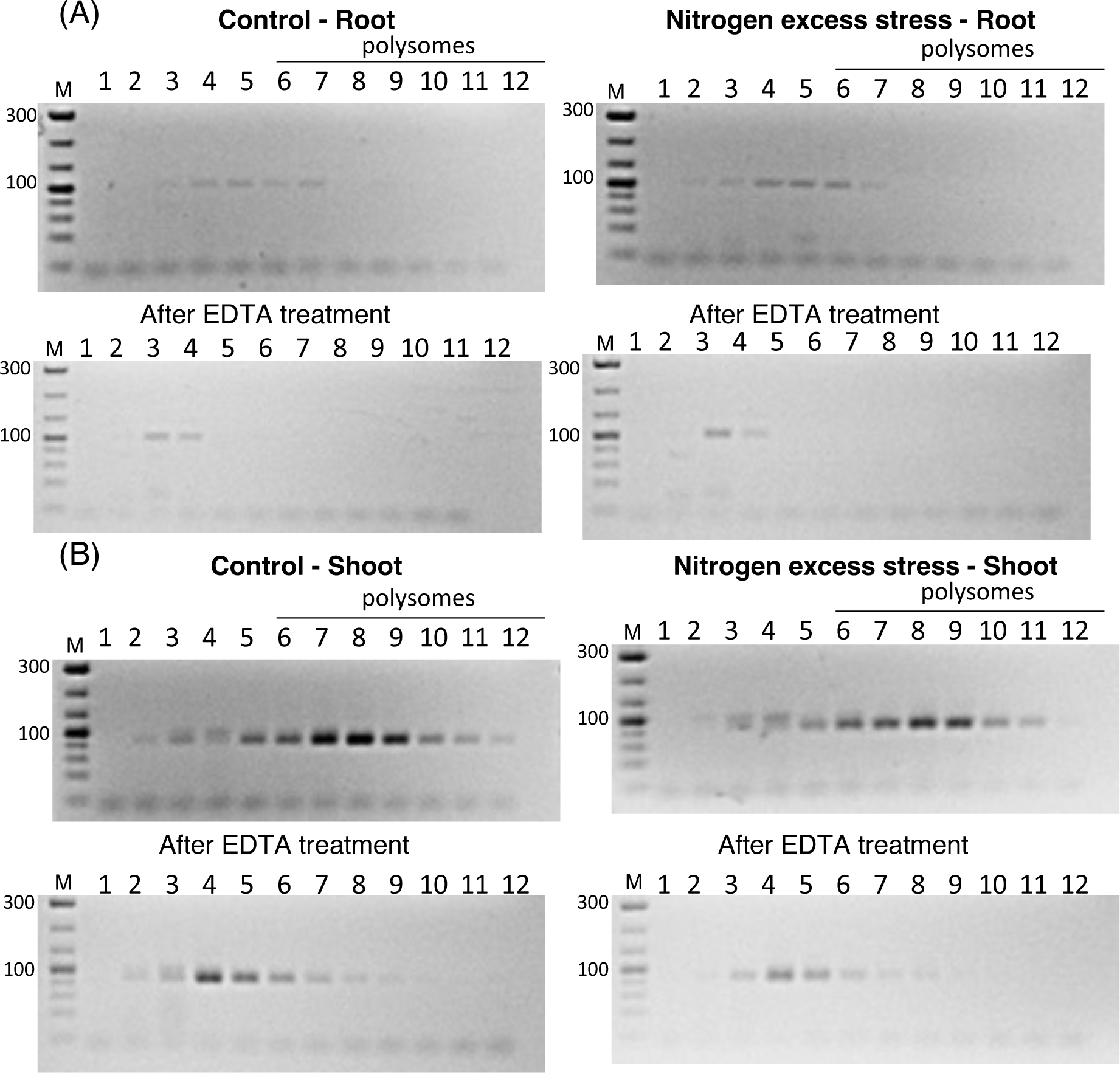
Polysome profiling of pri-miRNA444c extracted from barley roots and shoots grown under control and nitrogen excess stress conditions. RT-PCR analysis of the association of pri-miRNA444c with ribosomes in (A) roots and (B) shoots. Numbers given above the agarose gels correspond to the numbers of collected polysome profiling fractions. Polysomes as of the sixth fraction are marked. RT-PCR products observed of the sixth fraction indicate the specific binding of analysed pri-miRNA444c with ribosomes. After EDTA treatment, shift of RT-PCR products towards the fractions containing mRNA not associated to ribosomes is observed.. Primers used for pri-miRNA444c amplification encompassed the region encoding PEP444c. M-GeneRuler Low-Range DNA Ladder.

**Table 2.**
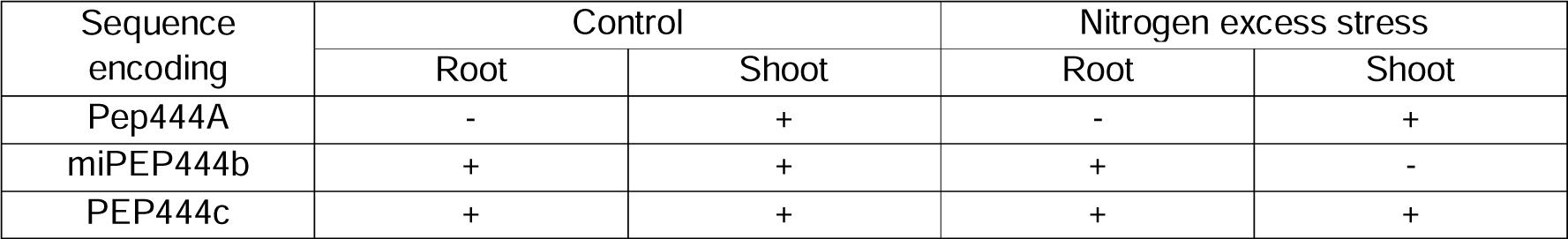
Summary of the association of sequences encoding PEP444a, miPEP444b, and PEP444c with ribosomes in shoot and root under control and excess nitrogen conditions, based on polysome profile absorbance and RT-PCR. Plus sign (+) indicates association with ribosomes, minus sign (-) depicts no association.

To confirm that PEP444a, miPEP444b, and PEP444c can be translated, overexpression of these peptides in *E. coli* was performed. All ORFs were fused to HisTaq and HaloTaq. To detect these peptides, rabbit custom-made antibodies raised against fragments of the analyzed peptides were used. The presence of PEP444a in soluble and insoluble fractions was detected. A weak signal was also observed in the control (in the absence of an inductor), which may suggest plasmid leakage (Golda et al., 2007). miPEP444b and PEP444c were detected mainly in soluble fractions after induction (Supplementary Figure 2).

Taken together, pri-miRNAs444 are exported to the cytoplasm, where they are associated with polysomes. This observation and overexpression experiments performed in *E. coli* support the idea that all studied ORFs may represent functional peptides in barley.

### *PEP444a* and *PEP444c* are expressed in barley

The presence of identified peptides in wild-type (WT) barley plants was tested. To detect these peptides, custom-made antibodies as described in the previous chapter were used. Serum collected prior to immunization of the rabbit was used to determine antibody specificity. PEP444a (13.7 kDa) in barley shoots was detected only upon nitrogen excess stress (Figure 12A, left panel). No specific signal was identified in the case of a Western blot performed with using serum sampled before immunization of the rabbit (Figure 12A, right panel). PEP444c (18.8 kDa) was identified in shoots and roots under control and excess nitrogen conditions (Figure 12B, left panel). As in the case of PEP444a, no specific signal was detectable on the Western performed using serum taken before immunization of the rabbit (Figure 12B, right panel). However, all attempts to detect miPEP444b were unsuccessful. Taken together, the presence of PEP444a and PEP444c in 2-week-old barley plants derived from *MIR444a* and *MIR444c* genes, was confirmed, respectively.

**Figure 12.**
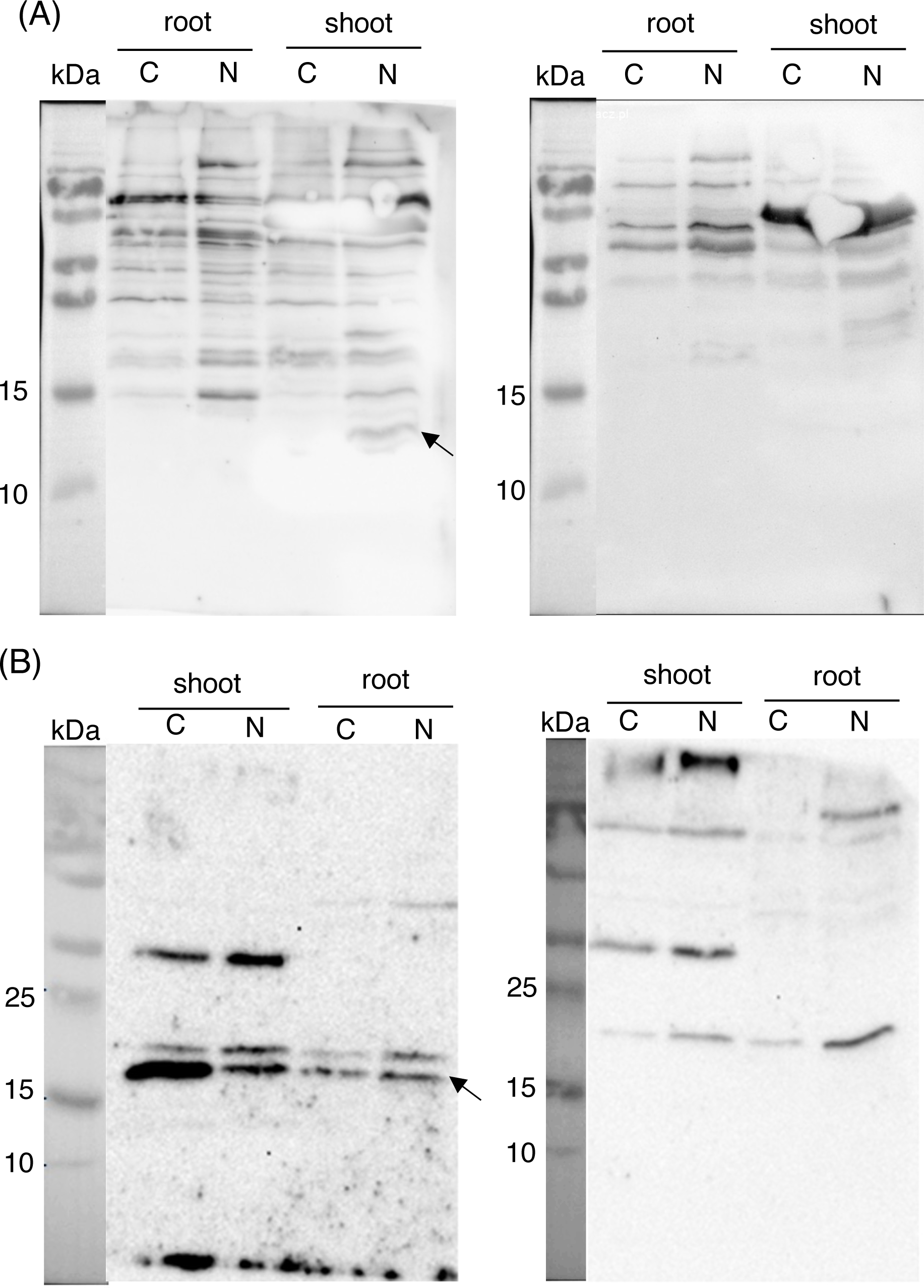
PEP444a and PEP444c are expressed in barley. (A) The left panel represents the detection of PEP444a in shoots upon nitrogen excess stress using Western blot and antibody against PEP444a. The right panel shows the membrane incubated with serum before immunization of the rabbit. (B) The left panel shows the detection of PEP444c in shoots and roots under control and nitrogen excess stress using Western blot and antibody against PEP444c. The right panel shows the membrane incubated with serum before immunization of the rabbit. (B) and (C) Arrows point to specific signals. C – control conditions; N – nitrogen excess stress; kDa – kilodaltons (PageRuler™ Prestained Protein Ladder Plus).

### Cas9-mediated generation of *pep444* mutants and characterization of transgenic plants

Based on the above results, the targeted mutagenesis approach was confined to PEP444a and PEP444c. Accordingly, gRNAs were designed for the sequences encoding both peptides (Supplementary Figure 3) and barley transformation was conducted using *cas9* and gRNA expression units. Analyses of T0 (primary mutant, M1) plants revealed that only plants were obtained without mutations within the sequence encoding PEP444a. This suggests that mutations in this region are lethal. However, mutations were found in the region of the *MIR444c* gene encoding PEP444c in M1 plants. Out of 60 analyzed M1 plants, nine proved to be mutated in *MIR444c* in homozygous condition as indicated by single peaks in the sequencing chromatograms.

M2 plants were analyzed to confirm inheritance of the mutations observed in M1. Line *pep444c-332.10* carrying a 1 bp insertion in the target motif of gRNA5 and a 9 bp deletion in the target motif of gRNA6, and line *pep444c-336.4/pep444c-336.6* carrying a 1 bp insertion in the target motif of gRNA5 and a 1 bp deletion in the target motif of gRNA6 were selected, respectively (Supplementary Figure 4). The mutations introduced within *MIR444c* affected the amino acid sequence of PEP444c. A premature stop codon introduced in PEP444c in *pep444c-332.10* transgenic plants (139 aa instead of 168 aa in WT) was identified, whereas, in *pep444c-336.4/pep444c-336.6* transgenic plants, 28 amino acids in PEP444c were modified owing to a shift in the translational reading frame (Figure 13A).

**Figure 13.**
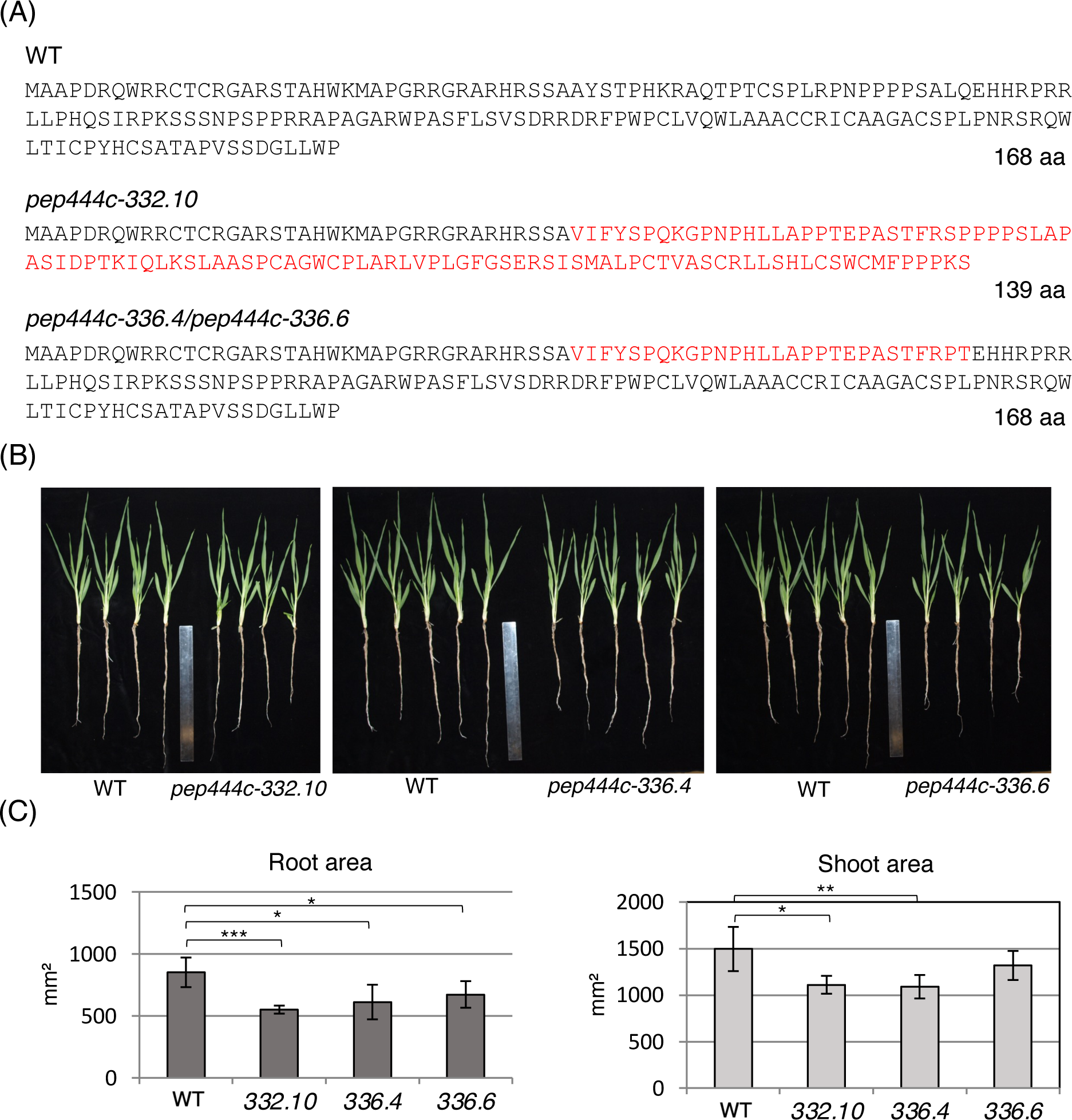
Characterization of *pep444c* mutant plants. (A) Amino acid sequences and length of PEP444c in WT, *pep444c-332.10, pep444c-336.4* and *pep444c-336.6* transgenic plants. (B) 2-week-old barley plants: WT, *pep444c-332.10, pep444c-336.4* and *pep444c-336.6.* (C) The measurements of root (left panel) and shoot (right panel) area of WT and mutant plants. Error bars indicate SD (n=5); asterisks indicate a significant difference between the sample and control (\**P* < 0.05; \*\*\**P* < 0.001).

Measurements of the shoot and root area in M2 plants of all three analyzed PEP444 mutant lines and WT plants were performed. There was a slight reduction of root (∼30%) and shoot (∼20%) surface area in mutated plants as compared to WT plants (Figure 13B, 13C). Chlorophyll fluorescence measurements were also performed. Analyzes of the quantum yield of regulatory non-photochemical quenching (NPQ), maximum fluorescence (Fm), minimum fluorescence (F0), fluorescence decrease ratio (Rfd), and maximum quantum efficiency of PSII photochemistry (Fv/Fm) did not show significant differences in mutant versus WT plants (Supplementary Figure 5). These data suggest that PEP444c mutations affect general aspects of plant development but do not affect photosynthetic performance.

### PEP444c regulates the accumulation of microRNA444c

The level of mutated *PEP444c* transcripts was measured. RT-qPCR analysis was carried out in the pri-miRNA444c region encompassing PEP444c. We observed a 5-fold downregulation of the transcript level in the mutants as compared to WT plants (Supplementary Figure 6).

To investigate whether mutations introduced in the PEP444c coding region impact the accumulation of corresponding microRNA, we performed small RNA sequencing using Illumina technology. We observed a decreased level of microRNA444c in roots of *pep444c-332.10* compared to WT plants, whereas the level of microRNA444c in shoots was unchanged (Figure 14). These results indicate a significant effect of PEP444c on the accumulation of microRNA444c in roots.

**Figure 14.**
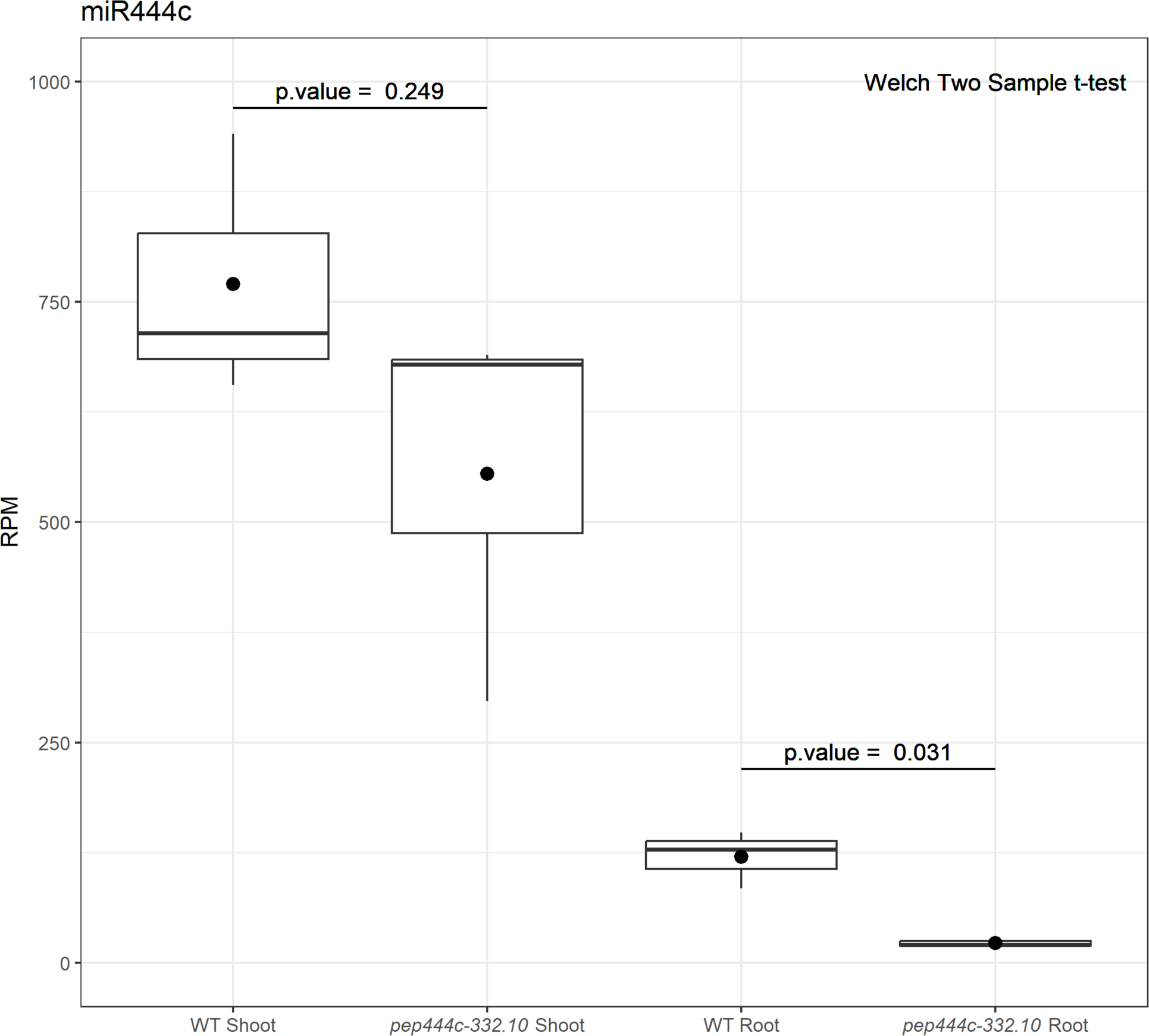
PEP444c regulates the level of microRNA444c. Analysis of microRNA444c levels based on small RNA sequencing results in shoots and roots of barley WT and *pep444c-332.10* mutant plants. The number of normalized counts for three biological replicates (RPM, reads per million) is shown. Welch’s t-test was used, p value indicates the significance level. For shoots, a significance level of 0.249 was noted (P > 0.05; not statistically significant), and for roots: 0.031 (P < 0.05; statistically significant).

Previous studies revealed that on the DNA strand opposite to the one containing *MIR444c*, the *MADS27* gene encoding a transcription factor from the MADS-box family is located. It was shown that MADS27 mRNA is not regulated by microRNA444c at least in two-week-old barley plants under control and excess nitrogen conditions. It has been also documented that the MADS27 transcription factor affects barley root architecture (Smoczynska et al., 2022). The *MADS27* gene does not overlap with the first exon of *MIR444c* gene. To verify that the mutations introduced within the *MIR444c* gene have no impact on *MADS27* expression, we analyzed the level of *MADS27* mRNA in plants derived from all three transgenic barley plants. RT-qPCR analyses of *MADS27* mRNA level did not reveal significant differences between analyzed mutants and WT plants (Supplementary Figure 7).

## DISCUSSION

In monocots, microRNAs from the *MIR444* family regulate the level of certain MADS-box transcription factors and thus play essential roles in plant growth, development, and response to environmental cues (Schilling et al., 2018; Schilling et al., 2020; Kuijer et al., 2021). *MIR444* genes in monocots are largely unexplored, which is why the detailed structure of these genes and the expression pattern of pri-miRNAs444 are still poorly understood. In the present study, three *MIR444* genes and their primary transcripts were comprehensively analysed in barley. It was found that the stress-mediated CREs were abundant in promoter regions of all of these genes. One of the most common CREs was CACTFTPPCA1 that is involved in the regulation of nitrogen metabolism in plants. Our earlier studies had revealed that the *HvMADS27* genomic sequence is localized at the opposite DNA strand of the *MIR444c* gene, and is strongly downregulated under nitrogen excess stress conditions. However, microRNA444c was not responsive to excess nitrogen in both shoots and roots (Smoczynska et al., 2022). Here, we analyzed the level of pri-miRNA444c and found that it is unchanged in roots upon nitrogen excess stress. Interestingly, we could not detect pri-miRNA444c in shoots under control and nitrogen excess stress conditions. The lack of correlation between the levels of pri-miRNA and mature microRNA has been previously described in *A. thaliana* and barley (Barciszewska-Pacak et al., 2015; Kruszka et al., 2013). Since microRNA444c is present in shoots, we can not exclude the possibility that it is transported from shoots to roots. The transport of 82 microRNAs from the shoot to the root and 6 microRNAs from the root to the shoot was previously demonstrated (Deng et al., 2021). Only pri-miRNA444b was slightly responsive to excess nitrogen stress, with its level being upregulated. However, the level of microRNA444b remained unchanged (Figure 1). Our results may reflect the complex regulation of microRNA biogenesis, which is thought to be influenced by splicing, alternative splicing, and the utilization of alternative polyadenylation sites (pA) (Bielewicz et al., 2013; Yan et al., 2012).

Barley pri-miRNA444a/b/c exhibit unusually intensive processing of pri-miRNA transcripts. We identified eight isoforms in the case of pri-miRNA444a as well as five in the case of pri-miRNA444b and pri-miRNA444c, respectively. Moreover, these isoforms can be divided into two groups: functional ones, in which the intron that separates exons encoding microRNA444 and miRNA444* is correctly spliced, and non-functional isoforms, in which one or both exons encoding microRNA444/444* are removed because of alternative splicing events. This unusual observation of the presence of functional and non-functional pri-miRNA444 isoforms may suggest an additional layer of microRNA444 level regulation via pri-miRNA splicing. It is known that environmentally induced alternative splicing is involved in the regulation of microRNA400 and microRNA402 in *A. thaliana* (Yan et al., 2012; Knop et al., 2017). Further studies are required to investigate this phenomenon in more detail. For instance, one of the identified isoforms of pri-miRNA444a (pri-miRNA444a.7, non-functional isoform) has been identified exclusively under nitrogen excess condition providing a possible mechanism of fine-tuning the level of microRNA444a.

Barley pri-miRNAs444 also undergo alternative polyadenylation (APA). In the case of pri-miRNA444c, APA sites were found in the first, second, and third intron. Since microRNA444c and microRNA444c* are encoded in the second and third exons, premature APA within the first and second intron may prevent microRNA444c production. Therefore, it seems plausible that the observed alternative polyadenylation sites in pri-miR444a/b/c may be of functional importance. Indeed, when we amplified pri-miRNA444c in the region that encompasses the first exon containing the PEP444c coding sequence, we detected its presence in shoots, while a simultaneously amplified pri-miRNA444c including the stem-loop structure was not detectable. Literature data indicate that more than 75% of genes in *Arabidopsis* undergo alternative polyadenylation (Guo et al., 2016). It has been shown that the choice of polyadenylation site can affect mRNA localization, stability, and translation efficiency (Wu et al., 2011; Guo et al., 2016). Earlier studies on pri-miRNA processing in plants also revealed the existence of many alternative polyadenylation sites. In the case of *Arabidopsis* microRNA402, high temperature inhibits splicing and activates a proximal polyadenylation site within the first intron, which entails increased accumulation of mature microRNA (Szarzynska et al., 2009; Knop et al., 2017).

It has been demonstrated that many pri-miRNAs contain ORFs encoding peptides, which in turn stimulates the accumulation of their corresponding microRNAs (Lauressergues et al., 2015; Couzigou and Combier, 2016; Chen et al., 2020, Sharma et al., 2020; Zhang et al., 2020). Our studies revealed that pri-miRNAs444 are associated with ribosomes. This experiment point to the presence of pri-miRNAs444 in the cytoplasm. Moreover, we provide experimental evidence that at least two identified peptides, PEP444a encoded by *MIR444a* and PEP444c encoded by *MIR444c*, are expressed in barley plants. Generally, we observed a correlation between ribosome profiling data and Western analyses. PEP444a and PEP444c do not significantly resemble any other peptides/proteins deposited in the NCBI and Ensembl databases. Therefore, our data indicate that peptides encoded by *MIR444* genes may not be present in other plant species, which is consistent with the literature data, according to which only three peptides encoded by the *MIR* genes (miPEP156a; miPEP164a and miPEP165a) identified so far are evolutionarily conserved in the *Brassicaceae* family (Lauressergues et al., 2015; Morozov et al., 2019; Lauressergues et al., 2022).

Generated transgenic lines with Cas9-mediated disruption of the PEP444c coding sequence (*pep444c-332.10*; *pep444c-336.4; pep444c-336.6*) show a decreased level of *PEP444c* transcript in shoots and roots. Moreover, mature microRNA444c is reduced five-fold in roots of *pep444c-332.10* mutant plants. Therefore, our data point to a significant effect of PEP444c on microRNA444c accumulation in roots. The parallel decrease of pri-miRNA444c and microRNA444c in roots may suggest the involvement of PEP444c in the early steps of microRNA444c biogenesis. Surprisingly, in shoots of *pep444c-332.10* mutant plants, we did not observe any effect on microRNA444c accumulation. This may be explained by the fact that we also did not observe pri-miRNA444c in shoots. Further experiments are required to clarify this phenomenon.

Our results revealed an additional layer of pri-miRNAs444 processing in barley. PEP444c, encoded by *MIR444c*, significantly affects the accumulation of microRNA444c in barley roots. The detailed mechanism by which pri-miRNA is directed into the cytoplasm and what is the mechanism by which PEP444c affects microRNA444c biogenesis is still unknown. In Figure 15, a model is presented showing the possible involvement of PEP444c in microRNA444 biogenesis.

**Figure 15.**
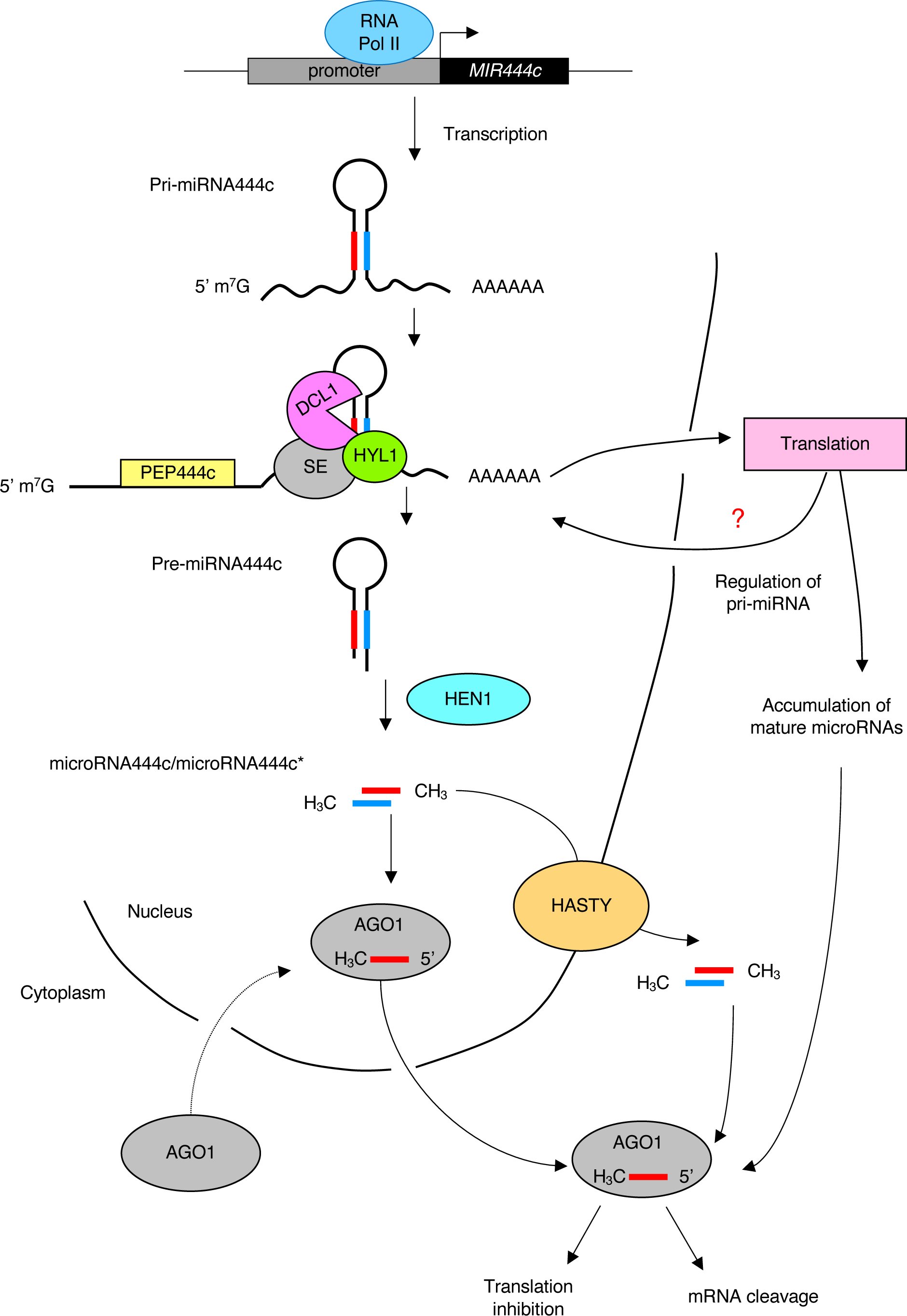
A model showing a possible effect of PEP444c encoded by the *MIR444c* gene in microRNA444c biogenesis. RNA Pol II – RNA Polymerase II; AAAAAA – poly(A) tail; DCL1 – DICER LIKE 1; SE – SERRATE; HYL1 - HYPONASTIC LEAVES 1; HEN1 – HUA ENHANCER 1, CH_3_ – methyl group; AGO1 - ARGONAUTE 1.

## MATERIALS AND METHODS

### Plant material and growth conditions

Grains of spring barley (*Hordeum vulgare* cultivar Golden Promise) were grown as described in Smoczynska et al. (2022). Briefly, grains were germinated under a 16/8h light/dark photoperiod at 20 °C until hypocotyls were visible. The seedlings were then transferred to pots containing perlite supplemented with medium as described by Pacak et al., 2016. To perform nitrogen excess stress, the concentration of NH_4_NO_3_ in the medium was increased to 280 mM, as described previously in Smoczynska et al. (2022). The plants were subsequently grown under controlled conditions (20 °C/15 °C day/night temperature) and 16/8 h light/dark photoperiod.

### Generation of constructs with *cas9* and gRNAs and plant transformation

Twenty nucleotide (nt) target sequences present immediately adjacent to a Protospacer Adjacent Motif (PAM) were selected using DESKGEN^TM^ CRISPR Libraries and verified using the RNAfold web server (http://rna.tbi.univie.ac.at/cgi-bin/RNAWebSuite/RNAfold.cgi) according to Kumlehn et al. (2018) for *MIR444a*, *MIR444b,* and *MIR444c* genes. Two single guide RNAs (gRNAs) for *MIR444a* (target-specific parts of gRNA1: AAGACATGCCCCTTGGTAGC and of gRNA2: ACAAAAGTGCCAGGCAGCTT), two gRNAs for *MIR444b* (target-specific parts of gRNA3: AGCCGGGAGTGCTTGGTCTT and of gRNA4: AAACCCTCGGCGCATGCAGG) and two gRNAs for *MIR444c* (target-specific parts of gRNA5: GTGGGGAGTAGAATATGCGG and of gRNA6: GGGGCGGTGGTGCTCTTGTA) were selected. The constructs targeting *MIR444a* (two guides) or *MIR444b/MIR444c* (four guides) were generated using the CasCADE modular vector system which is based on a hierarchical Golden Gate cloning strategy (Hoffie et al., in prep). Oligos (Table S3) with complementary target sequences and overhangs for cloning into BsaI restriction sites were annealed and cloned into BsaI– linearized vectors pIK1 to pIK4 containing the *OsU3* promoter. The resultant two or four vectors containing two/four gRNA units were digested with Esp3I enzyme and assembled into one vector, pIK195 (two guides) or pIK19 (four guides). To combine the multiple gRNA fragments with the *cas9* expression unit (pIK83) and an auxiliary unit vector (pIK155) into vector pIK22, BsaI restriction digest followed by ligation was used. Last, all expression units were mobilized into binary vector p6-d35S-TE9 (DNA Cloning Service) via SfiI restriction digest. The vectors containing *MIR444a*- or *MIR444b*/*MIR444c*-specific gRNAs were electroporated (Gene Pulser, Bio-Rad) into *Agrobacterium tumefaciens* strain AGL1 (Lazo et al., 1991) and DNA transfer into immature embryos of spring barley Golden Promise was performed (Hensel et al. 2008). Immature embryos were excised from caryopses 12–16 days after pollination and cultivated with *Agrobacterium* strain AGL1 carrying the respective vector for 48–72 h. Then, the explants were cultivated for callus induction under selective conditions using Timentin and hygromycin, followed by plant regeneration. DNA was extracted from leaves using Phire Plant Direct PCR Master Mix (Thermo Fischer Scientific), according to the manufacturer’s instructions. The target region was amplified by PCR and analyzed by Sanger sequencing using specific primers (Table S3).

### Phenotypic analysis of transgenic lines

The 2-week-old WT and *pep444c-332.10*, *pep444c-336.4* and *pep444c-336.6* mutant plants were automatically phenotyped for morphological traits and photosynthetic performance using the PlantScreen^TM^ Phenotyping System (Plant Photon Systems Instruments, Czech Republic). After 20 minutes of dark adaptation of the analyzed plants, chlorophyll *a* fluorescence and RGB imaging were conducted using a *Quenching* protocol pre-designed by manufacturer and a Plant Mask formula with modifications as described in Arasimowicz-Jelonek et al. (2022). Pixel count and fluorescence parameters from shoots were automatically evaluated from the obtained images. Root area was calculated separately after establishing an accurate Plant Mask formula.

### RNA isolation and quantitative PCR

According to the manufacturer’s instructions, total RNA was extracted from the shoots and roots using the Direct-zol RNA Mini Prep Kit (Zymo Research). RNA quality and quantity were measured using a NanoDrop ND-1000 spectrophotometer. RNA integrity was estimated on 1.2% agarose gel. Three μg of DNA-free RNA was reverse-transcribed to generate cDNA using SuperScript III Reverse Transcriptase (Invitrogen, Carlsbad, CA, United States) and oligo(dT)18 (Novazym, Poland) primers. RT-qPCR was performed using Power SYBR® Green PCR Master MIX (Applied Biosystems, Warrington, United Kingdom) and two specific primers (final concentration 500 nM each) on an Applied Biosystem QuantStudio7 Flex Real-Time PCR System in 10 μl reaction volumes in 384-well plates. The following thermal profile parameters were used: 10 min at 95 °C, 40 cycles for 15 s at 95 °C, and 1 min at 60 °C. Each RT-qPCR reaction was performed independently for the three biological replicates. The barley *ARF1* (*ADP-RIBOSYLATION FACTOR 1-like*; GenBank: AJ508228.2) gene fragment was simultaneously amplified and detected as a reference. Expression levels were calculated with the relative quantification method (2^-ΔCt^) as fold-change value and presented in the form of log_10_2^-ΔCt^. Primers are listed in Supplementary Table 3.

### RACE analysis

The 5’ and 3’ experiments were conducted with the use of SMARTer® RACE cDNA Amplification Kit (Takara Bio, United States, Cat# 634860) and Advantage Polymerase as described by Kruszka et al. 2014. PCR products were cloned into the pGEM T-Easy vector (Promega) and sequenced (Faculty’s Laboratory of Molecular Biology Techniques, Adam Mickiewicz University in Poznan, Poland).

### Small RNA library preparation

Small RNA sequencing data of WT barley plant shoots and roots in the control and nitrogen excess stress used in this study were previously published by Grabowska et al. (2020) and deposited in the GO database under accession number GSE145799. Small RNA libraries from the shoots and roots of *pep444c-332.10* plants were prepared as described in Grabowska et al. (2020). Samples were sequenced on an Illumina platform (Genesupport SA, Switzerland). Obtained data are available under accession number GSE229727.

### Polysome profiling

Polysome profiling from barley shoots and roots was optimized and performed according to the protocol described by Mustroph et al., 2009. The 2-week-old barley shoots and roots were ground separately in liquid nitrogen. 1.2 g of powdered shoots and 2 g of powdered roots were placed in a 50 ml Falcon tube, and two volumes of polysome extraction buffer were added according to Mustroph et al. (2009). The samples were centrifuged for 30 minutes at 10,000 ×*g* at 4 °C. Next, the supernatant was transferred to a new 50 ml Falcon tube. 700 μl of obtained extract was applied to a chilled sucrose density gradient and centrifuged at 4 °C for 2 hours 45 minutes at 151,000 ×*g* using the SW 41 Ti rotor (Beckman Culture Life Sciences). After centrifugation, gradient fractions were collected from top to bottom while measuring the absorbance at 254 nm. Fractions were frozen in liquid nitrogen and stored at -80 °C.

### Overexpression of PEP444a, miPEP444b and PEP444c in Escherichia coli

Sequences encoding *PEP444a*, *miPEP444b,* and *PEP444c* were amplified using a cDNA template obtained from 2-week-old barley shoots and roots and Q5® High-Fidelity DNA Polymerase (New England BioLabs) according to the manufacturer’s instructions. The primer sequences are listed in Table S3. PCR products were cloned into pH6HTN His6 HaloTaq®T7 (Promega) according to the manufacturer’s instructions and transformed into *E. coli*. IPTG (Isopropyl β-d-1-thiogalactopyranoside) was added to a final concentration of 0.4 mM to induce overexpression. Overexpression was carried out at 18 °C for 16 hours with shaking at 170 rpm and centrifuged at 4 °C for 15 minutes at 4,500 ×*g*. Bacterial cells were suspended in Sonic Buffer containing 5 mM DTT, 50 mM Tris-HCl pH 7.5, 200 mM NaCl, 1 mM EDTA, 0.1 % Triton X-100, 1× protease inhibitor complete EDTA-free (Roche), sonicated (15 cycles: 35 s ON, 35 s OFF) and centrifugated at 4 °C for 15 minutes at 14,000 ×*g*. The supernatant was collected (soluble fraction), and the pellet was suspended in Sonic Buffer (insoluble fraction).

### Western blot analysis

Western blot analysis was, in principle, performed as described in Smoczynska et al. (2022). Briefly, protein extracts were prepared using a buffer containing 50 mM Tris– HCl pH 7.5, 100 mM NaCl, 1 % Triton X-100, 1 mM EDTA, 10 mM NaF, 1 mM Na_3_VO_4_, 1% NP-40, 1 mM PMSF, 1×protease inhibitor complete EDTA-free (Roche), and 10 μM MG132. Protein extracts were separated on 13% SDS-PAGE, transferred to a polyvinylidene difluoride membrane (PVDF; Millipore), and analyzed by Western blotting using antibodies: anti-PEP444a (Agrisera, HORVU6), anti-miPEP444b (Agrisera, F2DYV0), anti-PEP444c (Agrisera, F2DAQ5), and anti-rabbit (Agrisera, AS09 602).

### Bioinformatic tools

The New PLACE database (A database of Plant Cis-acting Regulatory DNA Elements) (https://www.dna.affrc.go.jp/PLACE/?action=newplace) was used to analyze the regulatory elements of promoters of genes from the *MIR444* gene family. Genomic sequences for *MIR444a/b/c* were obtained from the Ensembl Plants database (https://plants.ensembl.org/Hordeum_vulgare/Info/Index) and aligned with the cDNA sequences using MAFFT version 7.4. An initial small RNA sequencing data quality analysis was performed using the Fastqc (version 0.11.5) program according to Wingett and Andrews (2018). The adapter sequences were removed using the cutadapt program (version 4.1) as described in Martin et al. (2011) and reads of 18 to 28 nucleotides in length were selected for further analysis. To count reads, the fastx_collapser function from the FASTX-toolkit (version 0.0.14) package was used. Further analyses (normalization of readings to RPM and statistical tests) were performed in the R programming environment. The ggplot2 package was used to visualize obtained results.

## Author contributions

AC performed the majority of the experiments included in this work and wrote the manuscript draft. AS took part in the analysis of promoter elements of *MIR444* genes and the nitrogen excess stress experiment. DB and WMK carried out bioinformatic analyses. AP participated in the analysis of the *MIR444* genes structure. GH and JK participated in the targeted mutagenesis approach. MG and ESN performed the phenotypic measurements of transgenic plants. AJ contributed to the discussion. ZS-K designed experiments and participated in manuscript writing. All authors contributed to the article and approved the submitted version.

## Funding

This study was funded by the PRELUDIUM project (2019/35/N/NZ9/01971) and OPUS project (2016/23/B/NZ9/00862) from the National Science Centre, Poland. Production of transgenic and mutant barley plants was possible thanks to the Visiting Programme for Scientists from Transition Countries in Europe and the former Soviet Union founded by the Leibniz Institute of Plant Genetics and Crop Plant Research (Gatersleben, Germany). Authors also received financial support from the Initiative of Excellence—Research University (05/IDUB/2019/94) at Adam Mickiewicz University, Poznan.

## Data availability statement

The original sRNA data are deposited and publicly available. These data can be found under accession number GSE229727.

## Acknowledgments

We thank Andrea Knospe and Sabine Sommerfeld for assisting in barley transformation.

## SUPPORTING INFORMATION

Additional supporting information can be found online in the Supporting Information section at the end of this article.

## Notes

### Competing Interest Statement

The authors have declared no competing interest.

## References

Arasimowicz-Jelonek, M., Jagodzik, P., Plociennik, A., Sobieszczuk-Nowicka, E., Mattoo, A., Polcyn, W., Floryszak-Wieczorek, J. (2022) Dynamics of nitration during dark-induced leaf senescence in Arabidopsis reveals proteins modified by tryptophan nitration. Journal of Experimental Botany, 73, 6853–6875. doi: 10.1093/jxb/erac341.

Bai, H., Euring, D., Volmer, K., Janz, D., Polle, A. (2013) The nitrate transporter (NRT) gene family in poplar. PLoS One, 19, 8, e72126. doi: 10.1371/journal.pone.0072126.

Barciszewska-Pacak, M., Milanowska, K., Knop, K., Bielewicz, D., Nuc, P., Plewka, P., Pacak, A.M., Vazquez, F., Karlowski, W., Jarmolowski, A., Szweykowska-Kulinska, Z. (2015) Arabidopsis microRNA expression regulation in a wide range of abiotic stress responses. Frontiers in Plant Science, 6, 410. doi: 10.3389/fpls.2015.00410.

Bartel, D. P. (2004). MicroRNAs: genomics, biogenesis, mechanism, and function. Cell, 116, 281–297. doi: 10.1016/S0092-8674(04)00045-5

Baumberger, N., Baulcombe, D.C. (2005) Arabidopsis ARGONAUTE1 is an RNA Slicer that selectively recruits microRNAs and short interfering RNAs. Proceedings of the National Academy of Sciences of the United States of America, 102, 11928–33. doi: 10.1073/pnas.0505461102.

Belity, T., Horowitz, M., Hoffman, J.R., Epstein, Y., Bruchim, Y., Todder. D., Cohen, H. (2022) Heat-Stress Preconditioning Attenuates Behavioral Responses to Psychological Stress: The Role of HSP-70 in Modulating Stress Responses. International Journal of Molecular Sciences, 23, 4129. doi: 10.3390/ijms23084129.

Bielewicz, D., Kalak, M., Kalyna, M., Windels, D., Barta, A., Vazquez, F., Szweykowska-Kulinska, Z., Jarmolowski, A. (2013) Introns of plant pri-miRNAs enhance miRNA biogenesis. EMBO Reports, 14, 622–8. doi: 10.1038/embor.2013.62.

Bologna, N.G., Iselin, R., Abriata, L.A., Sarazin, A., Pumplin, N., Jay, F., Grentzinger, T., Dal Peraro, M., Voinnet O. (2018) Nucleo-cytosolic Shuttling of ARGONAUTE1 Prompts a Revised Model of the Plant MicroRNA Pathway. Molecular Cell, 69, 709–719.e5. doi: 10.1016/j.molcel.2018.01.007.

Carlsbecker, A., Lee, J.Y., Roberts, C.J., Dettmer, J., Lehesranta, S., Zhou, J., Lindgren, O., Moreno-Risueno, M.A., Vatén, A., Thitamadee, S., Campilho, A., Sebastian, J., Bowman, J.L., Helariutta, Y., Benfey, P.N. (2010) Cell signalling by microRNA165/6 directs gene dose-dependent root cell fate. Nature, 465, 316–21. doi: 10.1038/nature08977.

Chen, Q.J., Deng, B.H., Gao, J., Zhao, Z.Y., Chen, Z.L., Song, S.R., Wang, L., Zhao, L.P., Xu, W.P., Zhang, C.X., Ma, C., Wang, S.P. (2020) A miRNA-Encoded Small Peptide, vvi-miPEP171d1, Regulates Adventitious Root Formation. Plant Physiology, 183, 656–670. doi: 10.1104/pp.20.00197.

Cheng, H., Hao, M., Wang, W., Mei, D., Tong, C., Wang, H., Liu, J., Fu, L., Hu, Q. (2016) Genomic identification, characterization and differential expression analysis of SBP-box gene family in Brassica napus. BMC Plant Biology, 16, 196.doi: 10.1186/s12870-016-0852-y.

Combier, J.P., Andre, O. (2018) Novel method for promoting nodulation in plants, in: U.S. Patent Application, 15/559,846.

Combier, J.P., Lauressergues, D., Becard, G., Payre, F., Plaza, S., Cavaille, J. (2016) Micropetides and use of same for modulating gene expression, U.S. Patent Application, 15/143,963.

Couzigou, J.M., Combier, J.P. (2016) Plant microRNAs: key regulators of root architecture and biotic interactions. New Phytologist, 212, 22–35. doi: 10.1111/nph.14058.

Deng, Z., Wu, H., Li, D., Li, L., Wang, Z., Yuan, W., Xing, Y., Li, C., Liang, D. (2021) Root-to-Shoot Long-Distance Mobile miRNAs Identified from Nicotiana Rootstocks. International Journal of Molecular Sciences, 22, 12821. doi: 10.3390/ijms222312821.

Grabowska, A., Bhat, S.S., Smoczynska, A., Bielewicz, D., Jarmolowski, A., Kulinska, Z.S. (2020) Regulation of Plant microRNA Biogenesis. In: Miguel, C., Dalmay, T., Chaves, I. (eds) Plant microRNAs. Concepts and Strategies in Plant Sciences. Springer International Publishing AG. https://doi.org/10.1007/978-3-030-35772-6_1

Guo, C., Spinelli, M., Liu, M., Li, Q.Q., Liang, C. (2016) A Genome-wide Study of “Non-3UTR” Polyadenylation Sites in Arabidopsis thaliana. Scientific Reports, 6, 28060. doi: 10.1038/srep28060.

Guo, S., Xu, Y., Liu, H., Mao, Z., Zhang, C., Ma, Y., Zhang, Q., Meng, Z., Chong, K. (2013) The interaction between OsMADS57 and OsTB1 modulates rice tillering via DWARF14. Nature Communications, 4, 1566. doi: 10.1038/ncomms2542.

Hensel, G., Valkov, V., Middlefell-Williams, J., Kumlehn, J. (2008) Efficient generation of transgenic barley: the way forward to modulate plant-microbe interactions. Journal of Plant Physiology, 165, 71–82. doi: 10.1016/j.jplph.2007.06.015.

Higo, K., Ugawa, Y., Iwamoto, M., Korenaga, T. (1999) Plant cis-acting regulatory DNA elements (PLACE) database: 1999. Nucleic Acids Research, 27, 297–300. doi: 10.1093/nar/27.1.297. PMID: 9847208; PMCID: PMC148163.

Hoffie, I., Daghma, D.E.S., Mirzakhmedov, M., Chamas, S., Egorova, A., Fontana, I.M., Hoffie, R.E., Ehrhardt, M., Marthe, C., Büchner, H., Hiekel, S., Kumlehn, J. CasCADE: A modular and versatile vector system for Cas endonuclease-mediated genome modifications validated in mono- and dicotyledonous plants. In preparation.

Hudson, M.E., Quail, P.H. (2003) Identification of promoter motifs involved in the network of phytochrome A-regulated gene expression by combined analysis of genomic sequence and microarray data. Plant Physiology, 133, 1605–16. doi: 10.1104/pp.103.030437.

Jia, F., Rock, C. D. (2013) MIR846 and MIR842 comprise a cistronic MIRNA pair that is regulated by abscisic acid by alternative splicing in roots of Arabidopsis. Plant Molecular Biology, 81, 447–460. doi: 10.1007/s11103-013-0015-6

Jiao, X., Wang, H., Yan, J., Kong, X., Liu, Y., Chu, J., Chen, X., Fang, R., Yan, Y. (2020) Promotion of BR Biosynthesis by miR444 Is Required for Ammonium-Triggered Inhibition of Root Growth. Plant Physiology, 182, 1454–1466. doi: 10.1104/pp.19.00190.

Knop, K., Stepien, A., Barciszewska-Pacak, M., Taube, M., Bielewicz, D., Michalak, M., Borst, J.W., Jarmolowski, A., Szweykowska-Kulinska, Z. (2017) Active 5’ splice sites regulate the biogenesis efficiency of Arabidopsis microRNAs derived from intron-containing genes. Nucleic Acids Research, 45, 2757–2775. doi: 10.1093/nar/gkw895.

Kruszka, K., Pacak, A., Swida-Barteczka, A., Nuc, P., Alaba, S., Wroblewska, Z., Karlowski, W., Jarmolowski, A., Szweykowska-Kulinska, Z. (2014) Transcriptionally and post-transcriptionally regulated microRNAs in heat stress response in barley. Journal of Experimental Botany, 65,6123–35. doi: 10.1093/jxb/eru353.

Kuijer, H.N.J., Shirley, N.J., Khor, S.F., Shi, J., Schwerdt, J., Zhang, D., Li, G., Burton, R.A. (2021) Transcript Profiling of MIKCc MADS-Box Genes Reveals Conserved and Novel Roles in Barley Inflorescence Development. Frontiers in Plant Science, 12, 705286. doi: 10.3389/fpls.2021.705286.

Kumlehn, J., Pietralla, J., Hensel, G., Pacher, M. Puchta, H. (2018) The CRISPR/Cas revolution continues: From efficient gene editing for crop breeding to plant synthetic biology. Journal of Integrative Plant Biology, 60, 1127–1153. doi: 10.1111/jipb.12734.

Kurihara, Y., Takashi, Y., and Watanabe, Y. (2006). The interaction between DCL1 and HYL1 is important for efficient and precise processing of pri-miRNA in plant microRNA biogenesis. RNA, 12, 206–212. doi: 10.1261/rna.2146906

Laubinger, S., Sachsenberg, T., Zeller, G., Busch, W., Lohmann, J. U., Ratsch, G., et al. (2008). Dual roles of the nuclear cap-binding complex and SERRATE in pre-mRNA splicing and micro RNA processing in Arabidopsis thaliana. Proceedings of the National Academy of Sciences of the United States of America, 8795–8800. doi: 10.1073/pnas.0802493105

Lauressergues, D., Couzigou, J.M., Clemente, H.S., Martinez, Y., Dunand, C., Bécard, G., Combier, J.P. (2015) Primary transcripts of microRNAs encode regulatory peptides. Nature, 520, 90–3. doi: 10.1038/nature14346.

Lauressergues, D., Ormancey, M., Guillotin, B., San Clemente, H., Camborde, L., Duboé, C., Tourneur, S., Charpentier, P., Barozet, A., Jauneau, A., Le Ru, A., Thuleau, P., Gervais, V., Plaza, S., Combier, J.P. (2022) Characterization of plant microRNA-encoded peptides (miPEPs) reveals molecular mechanisms from the translation to activity and specificity. Cell Reports, 38, 110339. doi: 10.1016/j.celrep.2022.110339.

Lazo, G.R., Stein, P.A., Ludwig, R.A. (1991) A DNA transformation-competent Arabidopsis genomic library in Agrobacterium. Biotechnology (N Y), 9, 963–7. doi: 10.1038/nbt1091-963.

Li, J., Yang, Z., Yu, B., Liu, J., Chen, X. (2005) Methylation protects miRNAs and siRNAs from a 3’-end uridylation activity in Arabidopsis. Current Biology, 15, 1501–7. doi: 10.1016/j.cub.2005.07.029.

Lobbes, D., Rallapalli, G., Schmidt, D.D., Martin, C., Clarke, J. (2006) SERRATE: a new player on the plant microRNA scene. EMBO Reports, 7, 1052–8. doi: 10.1038/sj.embor.7400806.

Lu, C., Jeong, D.H., Kulkarni, K., Pillay, M., Nobuta, K., German, R., Thatcher, S.R., Maher, C., Zhang, L., Ware, D., Liu, B., Cao, X., Meyers, B.C. (2008) Green PJ. Genome-wide analysis for discovery of rice microRNAs reveals natural antisense microRNAs (nat-miRNAs). Proceedings of the National Academy of Sciences of the United States of America, 105, 4951–6. doi: 10.1073/pnas.0708743105.

Manavella, P. A., Hagmann, J., Ott, F., Laubinger, S., Franz, M., Macek, B., et al. (2012). Fast-forward genetics identifies plant CPL phosphatases as regulators of miRNA processing factor HYL1. Cell, 151, 859–870. doi: 10.1016/j.cell.2012.09.039.

Morozov, S.Y., Ryazantsev, D.Y., Erokhina, T.N. (2019) Bioinformatics Analysis of the Novel Conserved Micropeptides Encoded by the Plants of Family Brassicaceae. Journal of Bioinformatics and Systems Biology, 2, 066–077.

Mustroph, A., Juntawong, P., Bailey-Serres, J. (2009) Isolation of plant polysomal mRNA by differential centrifugation and ribosome immunopurification methods. Methods in Molecular Biology, 553, 109–26. doi: 10.1007/978-1-60327-563-7_6.

Nag, A., King, S., Jack, T. (2009) miR319a targeting of TCP4 is critical for petal growth and development in Arabidopsis. Proceedings of the National Academy of Sciences of the United States of America, 106, 22534–9. doi: 10.1073/pnas.0908718106.

Pacak, A. M., Barciszewska-Pacak, M., Swida-Barteczka, A., Krusza, K., Sega, P., Milanowska, K., et al. (2016). Heat stress affects Pi-related genes expression and inorganic phosphate deposition/accumulation in barley. Frontiers in Plant Science, 7, 926. doi: 10.3389/fpls.2016.00926

Reinhart, B.J., Weinstein, E.G., Rhoades, M.W., Bartel, B., Bartel, D.P. (2002) MicroRNAs in plants. Genes & Development, 16, 1616–26. doi: 10.1101/gad.1004402.

Ren, G., Xie, M., Dou, Y., Zhang, C., and Yu, B. (2012) Regulation of miRNA abundance by RNA binding protein TOUGH in Arabidopsis. Proceedings of the National Academy of Sciences of the United States of America, 109, 12817–12821. doi: 10.1073/pnas.1204915109

Rieping, M., Schöffl, F. (1992) Synergistic effect of upstream sequences, CCAAT box elements, and HSE sequences for enhanced expression of chimaeric heat shock genes in transgenic tobacco. Molecular & general genetics, 231, 226–32. doi: 10.1007/BF00279795.

Rogers, K., Chen, X. (2013) Biogenesis, turnover, and mode of action of plant microRNAs. Plant Cell, 25, 2383–99. doi: 10.1105/tpc.113.113159.

Schilling, S., Kennedy, A., Pan, S., Jermiin, L.S., Melzer, R. (2020) Genome-wide analysis of MIKC-type MADS-box genes in wheat: pervasive duplications, functional conservation and putative neofunctionalization. New Phytologist, 225, 511–529. doi: 10.1111/nph.16122.

Schilling, S., Pan, S., Kennedy, A., Melzer, R. (2018) MADS-box genes and crop domestication: the jack of all traits. Journal of Experimental Botany, 69, 1447–1469. doi: 10.1093/jxb/erx479.

Sharma, A., Badola, P.K., Bhatia, C., Sharma, D., Trivedi, P.K. (2020) Primary transcript of miR858 encodes regulatory peptide and controls flavonoid biosynthesis and development in Arabidopsis. Nature Plants, 6, 1262–1274. doi: 10.1038/s41477-020-00769-x.

Sharma, D., Tiwari, M., Pandey, A., Bhatia, C., Sharma, A., Trivedi, P.K. (2016) MicroRNA858 Is a Potential Regulator of Phenylpropanoid Pathway and Plant Development. Plant Physiology, 171, 944–59. doi: 10.1104/pp.15.01831.

Smoczynska, A., Pacak, A., Grabowska, A., Bielewicz, D., Zadworny, M., Singh, K., Dolata, J., Bajczyk, M., Nuc, P., Kesy, J., Wozniak, M., Ratajczak, I., Harwood, W., Karlowski, W.M., Jarmolowski, A., Szweykowska-Kulinska, Z. (2022) Excess nitrogen responsive HvMADS27 transcription factor controls barley root architecture by regulating abscisic acid level. Frontiers in Plant Science, 13, 950796. doi: 10.3389/fpls.2022.950796.

Stothard, P. (2000) The sequence manipulation suite: JavaScript programs for analyzing and formatting protein and DNA sequences. BioTechniques, 28, 1102, 1104. doi: 10.2144/00286ir01.

Sunkar, R., Girke, T., Jain, P.K., Zhu, J.K. (2005) Cloning and characterization of microRNAs from rice. Plant Cell, 17, 1397–411. doi: 10.1105/tpc.105.031682.

Sunkar, R., Jagadeeswaran, G. (2008) In silico identification of conserved microRNAs in large number of diverse plant species. BMC Plant Biology, 8, 37. doi: 10.1186/1471-2229-8-37.

Sunkar, R., Li, Y. F., and Jagadeeswaran, G. (2012). Functions of microRNAs in plant stress responses. Trends in Plant Science, 17, 196–203. doi: 10.1016/j.tplants.2012.01.010

Szarzynska, B., Sobkowiak, L., Pant, B.D., Balazadeh, S., Scheible, W.R., Mueller-Roeber, B., Jarmolowski, A., Szweykowska-Kulinska, Z. (2009) Gene structures and processing of Arabidopsis thaliana HYL1-dependent pri-miRNAs. Nucleic Acids Research, 37, 3083–93. doi: 10.1093/nar/gkp189.

Terzaghi, W.B., Cashmore, A.R. (1995) Light-regulated transcription. Annual Review of Plant Physiology and Plant Molecular Biology, 46, 445–74. doi: 10.1146/annurev.pp.46.060195.002305.

Tiwari, V., Patel, M.K., Chaturvedi, A.K., Mishra, A., Jha, B. (2016) Functional Characterization of the Tau Class Glutathione-S-Transferases Gene (SbGSTU) Promoter of Salicornia brachiata under Salinity and Osmotic Stress. PLoS One, 11, e0148494. doi: 10.1371/journal.pone.0148494.

Vaucheret, H., Vazquez, F., Crété, P., Bartel, D.P. (2004) The action of ARGONAUTE1 in the miRNA pathway and its regulation by the miRNA pathway are crucial for plant development. Genes & Development, 18, 1187–97. doi: 10.1101/gad.1201404.

Vazquez, F., Gasciolli, V., Crete, P., and Vaucheret, H. (2004) The nuclear dsRNA binding protein HYL1 is required for microRNA accumulation and plant development, but not posttranscriptional transgene silencing. Current Biology, 14, 346–351. doi: 10.1016/j.cub.2004.01.035

Villain, P., Mache, R., Zhou, D.X. (1996) The mechanism of GT element-mediated cell type-specific transcriptional control. Journal of Biological Chemistry, 271, 32593–8. doi: 10.1074/jbc.271.51.32593.

Voinnet, O. (2009) Origin, biogenesis, and activity of plant microRNAs. Cell, 136, 669–87. doi: 10.1016/j.cell.2009.01.046.

Wang, H., Jiao, X., Kong, X., Hamera, S., Wu, Y., Chen, X., Fang, R., Yan, Y. (2016) A Signaling Cascade from miR444 to RDR1 in Rice Antiviral RNA Silencing Pathway. Plant Physiology, 170, 2365–77. doi: 10.1104/pp.15.01283.

Wu, X., Liu, M., Downie, B., Liang, C., Ji, G., Li, Q.Q. (2011) Hunt AG. Genome-wide landscape of polyadenylation in Arabidopsis provides evidence for extensive alternative polyadenylation. Proceedings of the National Academy of Sciences of the United States of America, 108, 12533–8. doi: 10.1073/pnas.1019732108.

Yan, K., Liu, P., Wu, C.A., Yang, G.D., Xu, R., Guo, Q.H., Huang, J.G., Zheng, C.C. (2012) Stress-induced alternative splicing provides a mechanism for the regulation of microRNA processing in Arabidopsis thaliana. Molecular Cell, 48, 521–31. doi: 10.1016/j.molcel.2012.08.032.

Yan, Y., Wang, H., Hamera, S., Chen, X., Fang, R. (2014) miR444a has multiple functions in the rice nitrate-signaling pathway. The Plant Journal, 78, 44–55. doi: 10.1111/tpj.12446.

Yan, K., Liu, P., Wu, C.-A., Yang, G.-D., Xu, R., Guo, Q.-H., et al. (2012). Stress-induced alternative splicing provides a mechanism for the regulation of microRNA processing in Arabidopsis thaliana. Molecular Cell, 48, 521–531. doi: 10.1016/j.molcel.2012.08.032

Yang, S.W., Chen, H.Y., Yang, J., Machida, S., Chua, N.H., Yuan, Y.A. (2010) Structure of Arabidopsis HYPONASTIC LEAVES1 and its molecular implications for miRNA processing. Structure, 18, 594–605. doi: 10.1016/j.str.2010.02.006.

Yu, B., Yang, Z., Li, J., Minakhina, S., Yang, M., Padgett, R.W., Steward, R., Chen, X. (2005) Methylation as a crucial step in plant microRNA biogenesis. Science, 307, 932–5. doi: 10.1126/science.1107130.

Zhan, X., Wang, B., Li, H., Liu, R., Kalia, R. K., and Zhu, J. K. (2012). Arabidopsis proline-rich protein important for development and abiotic stress tolerance is involved in microRNA biogenesis. Proceedings of the National Academy of Sciences of the United States of America, 109, 18198–18203. doi: 10.1073/pnas.1216199109

Zhang, Q.L., Su, L.Y., Zhang, S.T., Xu, X.P., Chen, X.H., Li, X., Jiang, M.Q., Huang, S.Q., Chen, Y.K., Zhang, Z.H., Lai, Z.X., Lin, Y.L. (2020) Analyses of microRNA166 gene structure, expression, and function during the early stage of somatic embryogenesis in Dimocarpus longan Lour. Plant Physiology and Biochemistry, 147, 205–214. doi: 10.1016/j.plaphy.2019.12.014.

