## Supplementary figures for "PEP444c encoded within the *MIR444c* gene regulates microRNA444c accumulation in barley"

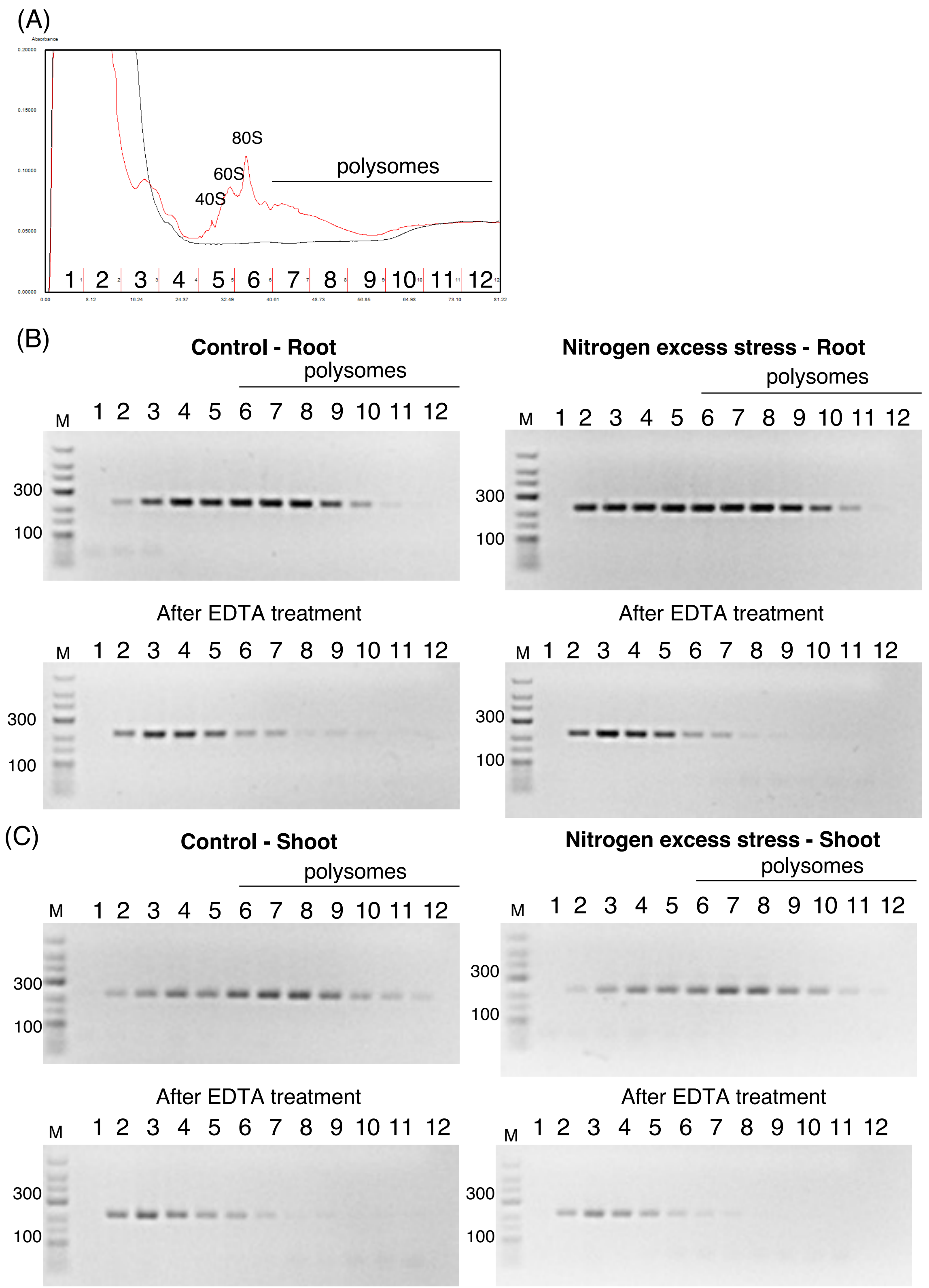


Figure S1. Polysome profiling of *ARF1* mRNA isolated from barley roots and shoots grown under control and nitrogen excess stress conditions. (A) An example of polysome profiling (red line) of mature ribosomes from barley shoots under control conditions. Black line represents absorbance profile after treatment with translation inhibitors, cycloheximide and chloramphenicol. RT-PCR analysis of the association of the *ARF1* mRNA with ribosomes in (B) roots and (C) shoots under control and nitrogen excess stress conditions. Numbers given above the agarose gels correspond to the numbers of the collected fractions shown in (A). Polysomes as of the sixth fraction are marked. RT-PCR products observed of the sixth fraction indicate the specific binding of analyzed pri-miRNAs444 with ribosomes. After EDTA treatment, shift of RT-PCR products towards the fractions containing mRNA not associated to ribosomes is observed. M- GeneRuler Low-Range DNA Ladder.

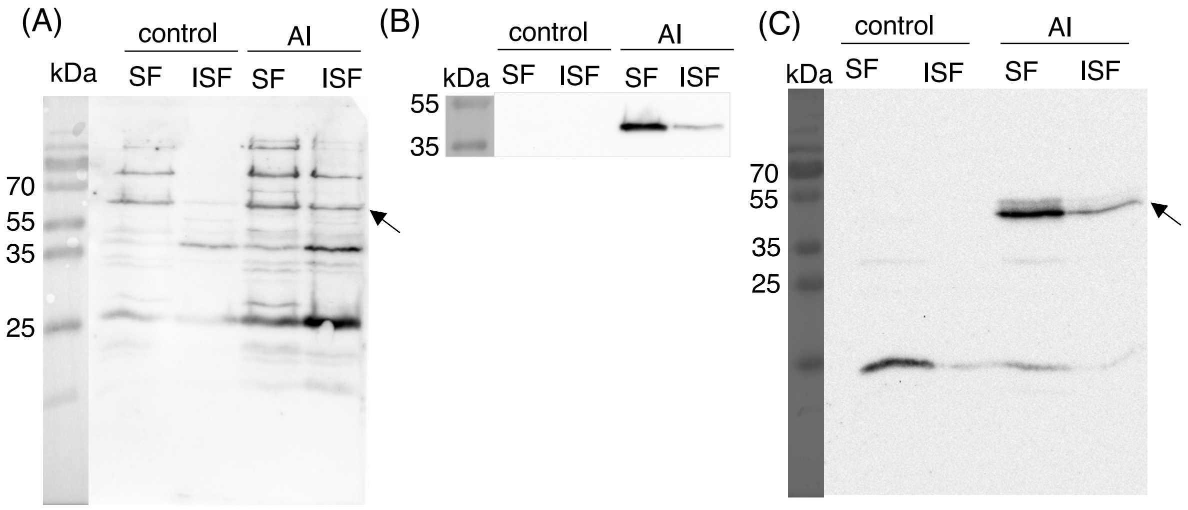


Figure S2. Peptides encoded by barley *MIR444a/b/c* genes are expressed in *E. coli*. Detection of PEP444a (A), miPEP444b (B), and PEP444c (C) using Western blot. Arrows indicate specific signals. SF – soluble fraction; ISF – insoluble fraction; AI – after induction; kDa – kilodaltons (PageRuler™ Prestained Protein Ladder Plus).

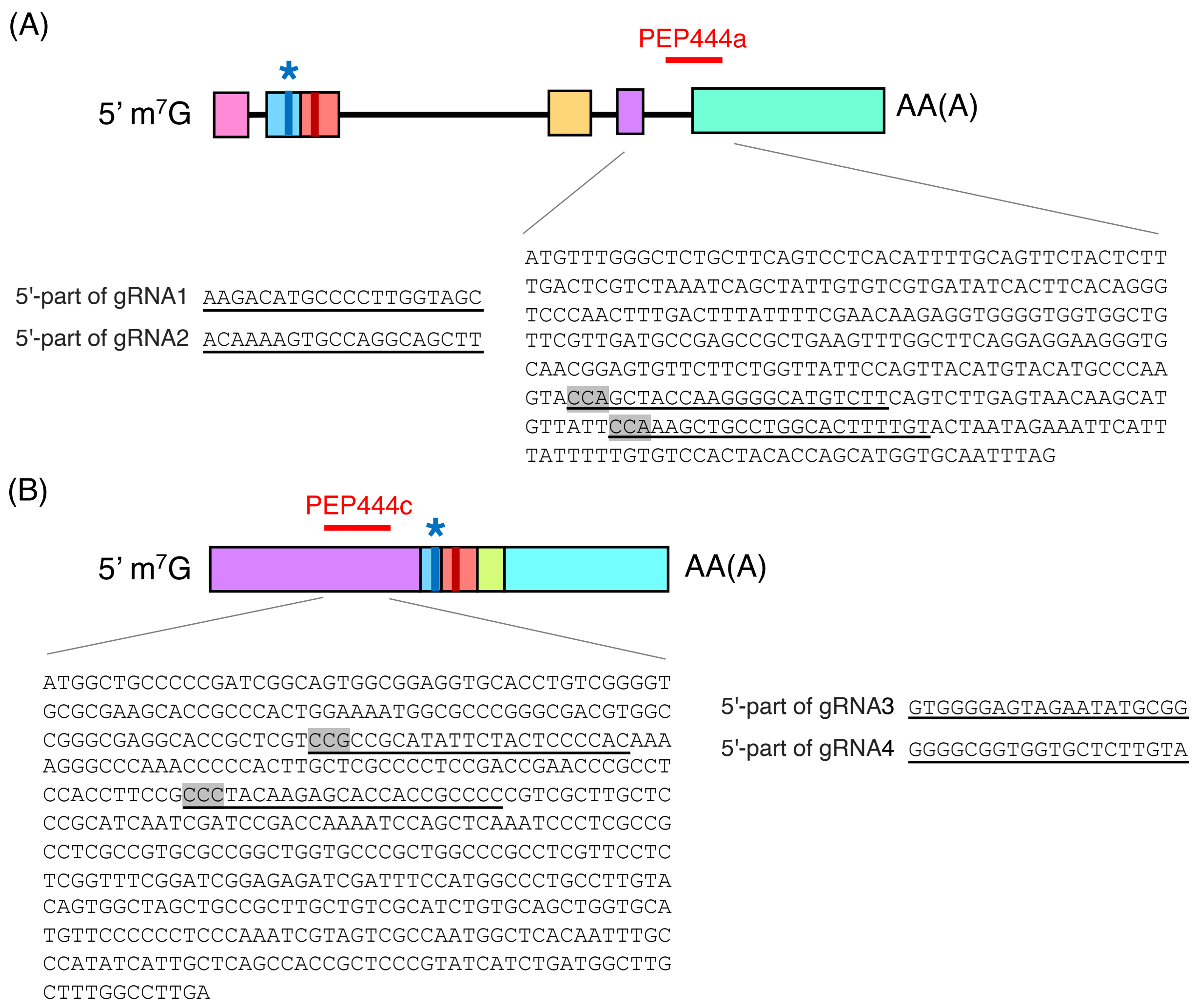


Figure S3. Construction of cas9/gRNA vectors targeting *MIR444a* and *MIR444b* gene areas encoding PEP444a and PEP444b, respectively. Precursors of *MIR444a* (A) and *MIR444c* (B) are presented. (A) and (B) Red lines indicate the positions of identified PEP444a and PEP444c. Nucleotide sequences encoding identified peptides and gRNAs are shown. The sequences of 5'-part of gRNAs (20 nt) are underlined, and PAM (NGG) sequences are highlighted in grey. Colored boxes represent exons, while black lines show introns. Blue bar and blue star represent microRNA* position, and red bar indicates microRNA position.

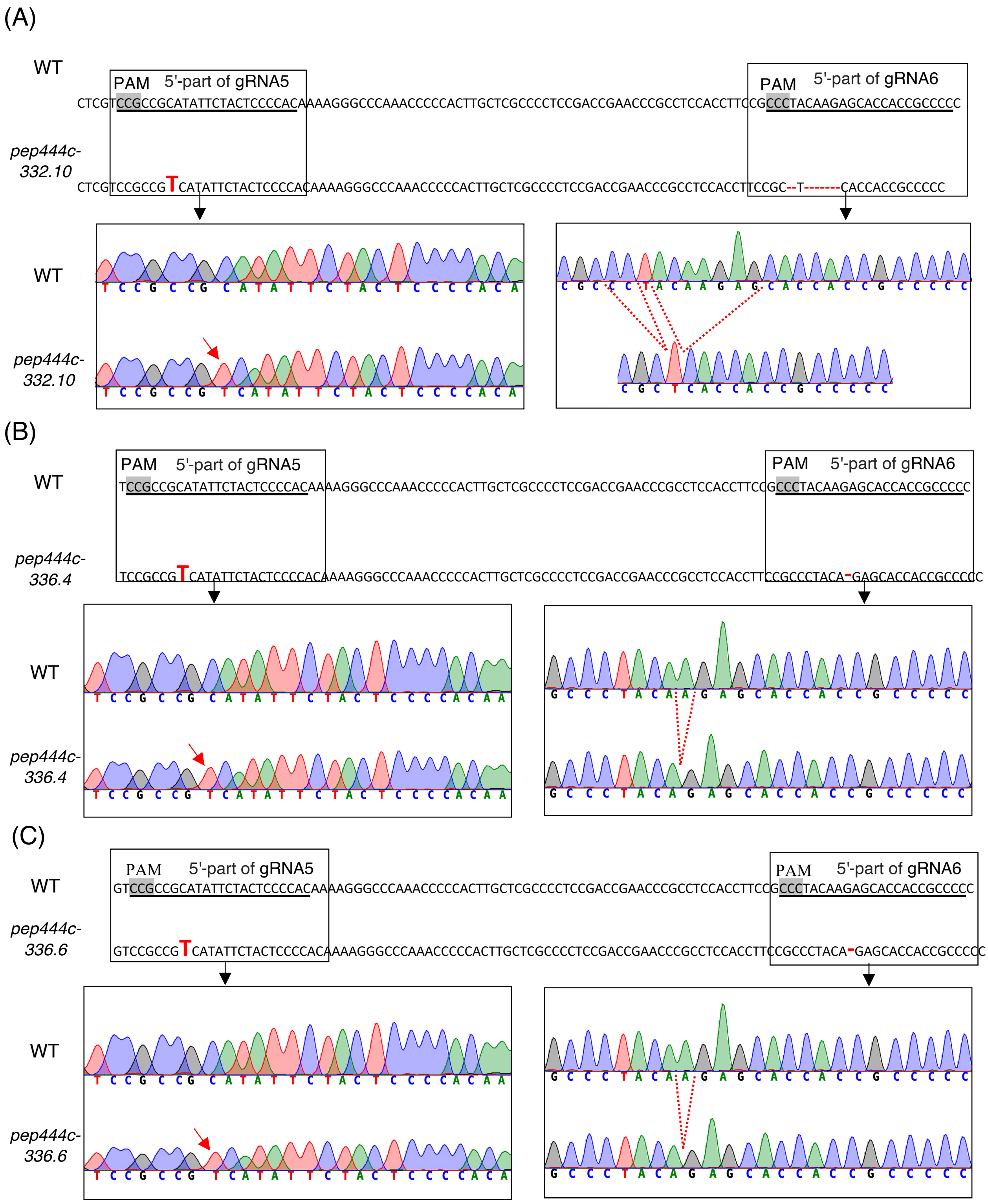


Figure S4. Cas9-induced mutation patterns in *pep444c -332.10* (A), *pep444c-336.4* (B), and *pep444c-336.6* (C) of barley transgenic plants. Target area of *MIR444c* in WT and homozygous mutations obtained in G2 *pep444c* mutants are shown. The PAM (NGG) sequence is highlighted in grey, the 20 bp long target region is underlined, and point mutations are enlarged, marked in red and bold. On chromatograms, insertions are marked with red arrows, and deletions are marked with red dotted lines. Below the nucleotide sequences, fragments of chromatograms show DNA sequences of selected *MIR444c* regions including the Cas9-induced mutations.

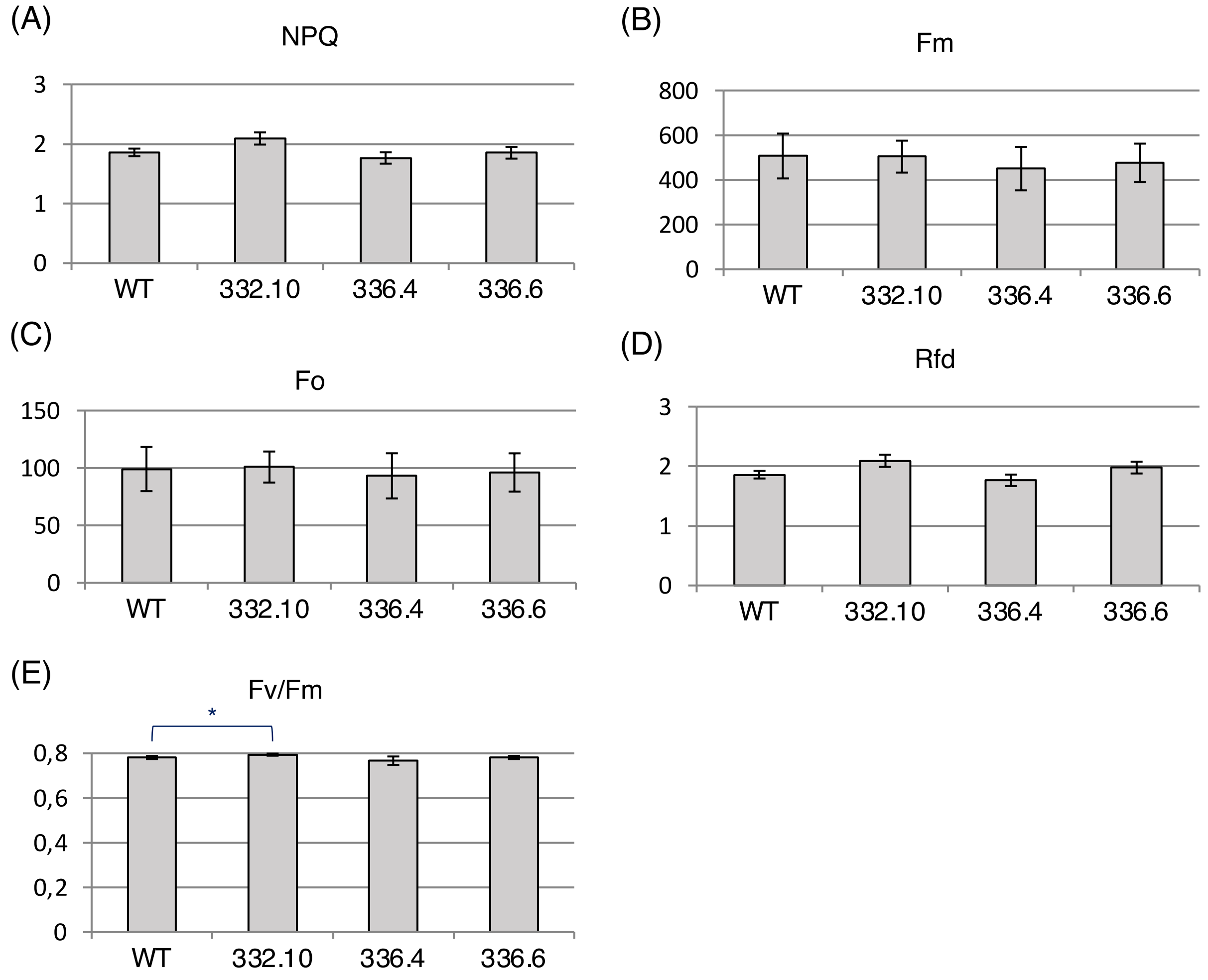


Figure S5. Chlorophyll fluorescence parameters in WT and *pep444c-332.10, pep444c-336.4,* and *pep444c-336.6* mutant plants. (A) The quantum yield of regulatory non-photochemical quenching (NPQ), (B) maximum fluorescence (Fm), (C) minimum fluorescence (F0), (D) fluorescence decrease ratio (Rfd), and (E) maximum quantum efficiency of PSII photochemistry (Fv/Fm) were obtained by applying the *Quenching* protocol to dark- adapted plants. Error bars indicate SD (n=5); an asterisk indicates a significant difference between samples and control (*$P < 0.05$).

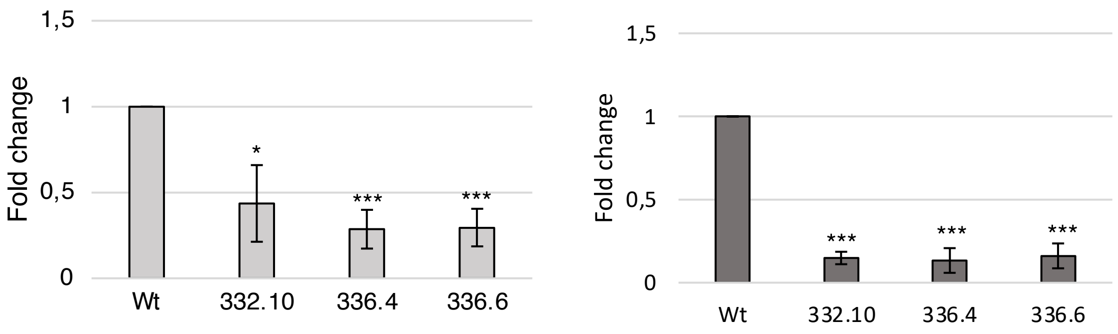


Figure S6. RT-qPCR analysis of PEP444c transcripts in shoots (left panel) and roots (right panel) in WT and transgenic barley plants derived from three independent Cas9-induced mutant lines. Error bars indicate SD (n=3); asterisks indicate a significant difference between samples and control (*$P < 0.05$; ***$P < 0.001$).

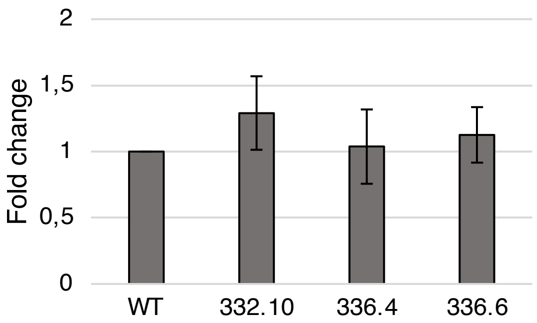


Figure S7. RT-qPCR analysis of *MADS27* mRNA in WT barley roots. Error bars indicate SD (n=3).
