## Supplementary tables for "PEP444c encoded within the *MIR444c* gene regulates microRNA444c accumulation in barley"

Table S1. The most abundant *cis*-regulatory elements identified in the promoters of *MIR444a/b/c* genes.

| **Motif** | **Sequence** | **Abundance in promoter regions of** | | | **Function** |
| --- | --- | --- | --- | --- | --- |
|  |  | ***MIR444a*** | ***MIR444b*** | ***MIR444c*** |  |
| CACTFTPPCA1 | YACT | 22 | 23 | 22 | Regulation of nitrogen metabolism in plants (Bai et al., 2013); Regulation of phosphoenolpyruvate carboxylase (Gowik et al., 2004) |
| ARR1AT | NGATT | 14 | 27 | 14 | Regulatory element involved in cytokinin response (Sakai et al., 2000) |
| EBOXBNNAPA | CANNTG | 7 | 26 | 20 | Regulatory element involved in light response (Tiwari et al., 2016) |
| MYCCONSENSUSAT | CANNTG | 7 | 26 | 20 | Regulatory element involved in dehydration response (Tiwari et al., 2016) |
| DOFCOREZM | AAAG | 12 | 21 | 17 | DOF transcription factor binding site (Yanagisawa and Schmidt, 1999) |
| CAATBOX1 | CAAT | 11 | 23 | 15 | Common in promoters of heat stress response genes (Rieping and Schöffl, 1992) |
| GT1CONSENSUS | GRWAAW | 12 | 24 | 6 | Regulatory element involved in light and salicylic acid response (Villain et al., 1996; Buchel et al., 1999) |
| CGCGBOXAT | VCGCGB | 14 | 4 | 18 | Regulatory element involved in auxin response (Yang and Poovaiah, 2002; Wong et al., 2017) |
| WRKY71OS | TGAC | 11 | 12 | 11 | WRKY71OS transcription binding site, regulation of gibberellin signaling pathway (Zhang et al., 2004) |
| GATABOX | GATA | 9 | 15 | 7 | Regulatory element involved in light response (Terzaghi and Cashmore, 1995) |
| CURECORECR | GTAC | 4 | 18 | 8 | Regulatory element involved in copper response (Tiwari et al., 2016) |
| CGACGOSAMY3 | CGACG | 10 | 3 | 15 | Common in promoters of genes encoding amylases (Hwang et al., 1998) |
| ACGTATERD1 | ACGT | 14 | 6 | 6 | Present in the promoter of the *ERD1* gene in *Arabidopsis* (Yamamoto et al., 2007) |
| GTGANTG10 | GTGA | 15 | 0 | 11 | Regulation of pollen-specific gene expression (Rogers et al., 2001) |
| POLLEN1LELAT52 | AGAAA | 5 | 13 | 5 | Involvement in the activation of the *LAT52* gene (Bate and Twell, 1998) |
| LTRECOREATCOR15 | CCGAC | 8 | 1 | 11 | Element involved in low-temperature response (Baker et al., 1994) |
| MYBCORE | CNGTTR | 5 | 4 | 11 | MYB protein binding site (Chen et al., 2019) |
| SORLIP1AT | GCCAC | 7 | 6 | 6 | Regulatory element involved in light response (Hudson and Quail, 2003) |
| PRECONSCRHSP70A | SCGAYNRNNNNNNNNNNNNNNNHD | 11 | 1 | 5 | Heat stress response element (Belity et al., 2022) |
| ROOTMOTIFTAPOX1 | ATATT | 5 | 8 | 2 | Involvement in the regulation of gene expression in root cells (Zhang et al., 2019) |
| BIHD1OS | TGTCA | 6 | 4 | 3 | BIHD1 transcription factor binding site (Liu et al., 2017) |
| RAV1AAT | CAACA | 4 | 6 | 3 | RAV1 transcription factor binding site (Hwang et al., 2008) |
| CCAATBOX1 | CCAAT | 2 | 6 | 4 | Heat stress response element (Rieping and Schöffl, 1992) |
| SEF4MOTIFGM7S | RTTTTTR | 2 | 8 | 2 | Regulatory element involved in embryo development (Silva et al., 2022) |
| WBOXNTERF3 | TGACY | 3 | 5 | 4 | Present in the promoter of the gene encoding ERF3 (Finkelstein et al., 2002) |
| CARGCW8GAT | CWWWWWWWWG | 3 | 6 | 2 | MADS transcription factor binding site (Aerts et al., 2018) |
| CBFHV | RYCGAC | 2 | 2 | 7 | Element involved in low-temperature response (Zarka et al., 2003) |
| DPBFCOREDCDC3 | ACACNNG | 5 | 2 | 4 | Abscisic acid (ABA) response element (Kim  et al., 1997) |
| MYBCOREATCYCB1 | AACGG | 1 | 2 | 8 | MYB transcription factor binding site; ABA response element (Abe et al., 2003) |
| MYB2CONSENSUSAT | YAACKG | 1 | 3 | 7 |  |
| SORLIP2AT | GGGCC | 2 | 0 | 9 | Regulatory element involved in light response (Hudson and Quail, 2003) |
| TATABOX5 | TTATTT | 1 | 9 | 1 | TATA sequence, binding site for the proteins involved in the process of transcription initiation (Patikoglou et al., 1999) |
| ABRERATCAL | MACGYGB | 5 | 3 | 2 | ABA response element (Yang et al., 2011; Kaur et al., 2017) |
| ABRELATERD1 | ACGTG | 8 | 1 | 1 |  |
| TAAAGSTKST1 | TAAAG | 2 | 5 | 3 | DOF transcription factor binding site (Plesch et al., 2001) |
| WBOXATNPR1 | TTGAC | 3 | 4 | 3 | WRKY transcription factor binding site (Yu et al., 2001) |
| CIACADIANLELHC | CAANNNNATC | 1 | 4 | 4 | Regulatory element involved in Circadian rhythm component (Piechulla et al., 1998) |
| GT1GMSCAM4 | GAAAAA | 2 | 5 | 2 | Element involved in salinity response (Liang et al., 2017) |
| IBOXCORE | GATAA | 1 | 7 | 1 | Regulatory element involved in light response (Li et al., 2014) |
| NODCON2GM | CTCTT | 1 | 4 | 4 | Involvement in the regulation of gene expression in root cells (Wang et al., 2022) |
| OSE2ROOTNODULE | CTCTT | 1 | 4 | 4 |  |
| ASF1MOTIFCAMV | TGACG | 2 | 2 | 4 | Auxins and salicylic acid response (Luo et al., 2013) |
| INRNTPSADB | YTCANTYY | 0 | 5 | 2 | Regulatory element involved in light response (Li et al., 2014) |
| POLASIG3 | AATAAT | 1 | 6 | 0 | Consensus sequence, polyadenylation signal (Joshi, 1987) |
| RHERPATEXPA7 | KCACGW | 2 | 2 | 3 | Element involved in the regulation of gene expression in root cells (Ahmadi et al., 2018) |
| TBOXATGAPB | ACTTTG | 3 | 2 | 2 | Regulatory element involved in light response (Yamamoto et al., 2007) |

Table S2. List of ORFs in pri-miRNAs444a/b/c. The peptides selected for experiments are marked in red.

| **pri-miRNA** | **Isoform** | **Number of ORFs** | **Sequence** | **The length of potential peptide** |
| --- | --- | --- | --- | --- |
| **pri-miRNA444a** | (A) | 6 | 1) MYRVLSGFGFKYFGENRITAVIILEILCIQDFWQSRDRTSCDVM 2) MYVPGLQICLKVKISHVWALLQSSHFAVLLFDSSKSAIVS 3) MELMVFLLSSRVVCNLYSLCKEIQKRLDQ 4) MLMVCRLARYACGGTKHEATTAVLVGKAQVGHQHYLQARRKINRTSPYLWLSCKSCSCCLKLAASLCQSYQKKT 5) MASSSICMIPLEERQHNFHFKSLPLSLRQLLPPIHITCRSLVQLLGSWRSRYTVTLPRWEK 6) MFGLCFSPHILQFYSLTRLNQLLCRDITSQGPNFDFIFEQEVGWWLFVDAEPLKFGFRRKGATECSSGYSSYMYMPKYQLPRGMSSVLSNKHVIPKLPGTFVLIEIHLFLCPLHQHGAI | 44 aa  40 aa  29 aa  74 aa  61 aa  119 aa |
|  | (B)  (C) | 4 | 1) MASSSICMIPLEERQHNFHFKSLPLSLRQLLPPIHITCRSLVQLLGSWRSRYTVTLPRWEK 2) MELMVFLLSSRVVCNLYSLCKEIQKRLDQ 3) MYRVLSGFGFKYFGENRITAVIILEILCIQDFWQSRDRTSCDVM 4) MYMPKYQLPRGMSSVLSNKHVIPKLPGTFVLIEIHLFLCPLHQHGAI | 61 aa  29 aa  44 aa  47 aa |
|  | (D) | 4 | 1) MYRVLSGFGFKYFGENRITAVIILEILCIQDFWQSRDRTSCDVM 2) MYMPKYQLPRGMSSVLSNKHVIPKLPGTFVLIEIHLFLCPLHQHGAI 3) MLMVCRLARYTCGGTKHEATTAVLVGKAQVGHQHYLQARRKINRTSPYLWLSCKSCSCCLKLAASLCQSYQKKT 4) MASSSICMIPLEERQHNFHFKSLPLSLRQLLPPIHITCRSLVQLLGSWRSRYTVTLPRWEK | 61 aa  29 aa  44 aa  47 aa |
|  | (E)  (H) | 2 | 1) MELMVFLLSSRVVCNLYSLCKEIQKRLDQ 2) MYMPKYQLPRGMSSVLSNKHVIPKLPGTFVLIEIHLFLCPLHQHGAI | 29 aa  47 aa |
|  | (F) | 3 | 1) MELMVFLLSSRVVCNLYSLCKEIQKRLDQ 2) MYMPKYQLPRGMSSVLSNKHVIPKLPGTFVLIEIHLFLCPLHQHGAI 3) MASSSICMIPLEERQHNFHFKSLPLSLRQLLPPIHITCRSLVQLLGSWRSRYTVTLPRWEK | 29 aa  47 aa  61 aa |
|  | (G) | 1 | 1) MYMPKYQLPRGMSSVLSNKHVIPKLPGTFVLIEIHLFLCPLHQHGAI | 47 aa |
| **pri-miRNA444b** | (A) | 6 | 1) MRRGFELLHVVAPSMRQQLHYFQGSYKIYGSS 2) MNDHTNKFEETFDSKLFTLVNTGNMDPWSISSPRHPSTP 3) MKPQIFVCCMWTKNKQINSCSRPSSCTLVVLF 4) MESSSARRSFTSRLLVLGPPLLPQPGVLGLGRAWPPACAEGLSYCMWWHQA 5) MGLHNRDFLASCAVAVSSLLTPSARPQCEIGNLVSFSE 6) MEHFFTASPLHTLKRTHLKASPASLKLTFTDILN | 32 aa  39 aa  32 aa  51 aa  38 aa  34 aa |
|  | (B) | 5 | 1) MRRGFELLHVVAPSMRQQLHYFQGSYKIYGSS 2) MEHFFTASPLHTLKRTHLKASPASLKLTFTDILS 3) MESSSARRSFTSRLLVLGPPLLPQPGVLGLGRAWPPACAEGLSYCMWWHQA 4) MGLHNRDFLASCAVAVSSLLTPSARPQCEIGNLVRRFCRMFLQSLNLELMCTQVNTGNMDPWSISSPRHPSTP 5) MKPQIFVCCMWTKNKQINSCSRPSSCTLVVLF | 32 aa  34 aa  51 aa  73 aa  32 aa |
|  | (C) | 5 | 1) MRRGFELLHVVAPSMRQQLHYFQGSYKIYGSS 2) MEHFFTASPLHTLKRTHLKASPASLKLTFTDILN 3) MESSSARRSFTSRLLVLGPPLLPQPGVLGLGRAWPPACAEGLSYCMWWHQA 4) MGLHNRDFLASCAVAVSSLLTPSARPQCEIGNLVRFFLLTSKSFFGCIKN 5) MKPQIFVCCMWTKNKQINSCSRPSSCTLVVLF | 32 aa  34 aa  51 aa  50 aa  32 aa |
|  | (D) | 5 | 1) MRRGFELLHVVAPSMRQQLHYFQGSYKIYGSS 2) MKPQIFVCCMWTKNKQINSCSRPSSCTLVVLF 3) MESSSARRSFTSRLLVLGPPLLPQPGVLGLGRAWPPACAEGLSYCMWWHQA 4) MGLHNRDFLASCAVAVSSLLTPSARPQCEIGNLVR 5) MEHFFTASPLHTLKRTHLKASPASLKLTFTDILN | 32 aa  32 aa  51 aa  35 aa  34 aa |
|  | (E) | 5 | 1) MKPQIFVCCMWTKNKQINSCSRPSSCTLVVLF 2) MESSSARRSFTSRLLVLGPPLLPQPGVLGLGRAWPPACAEGLR 3) MEHFFTASPLHTLKRTHLKASPASLKLTFTDILN | 32 aa  43 aa  34 aa |
| **pri-miRNA444c** | (A)  (C) | 3 | 1) MEFELTMSSGAKMPHSIHFMEGVKLSEHGRRRLGF 2) MVSYRSIPTTFVVSSWANICMVLRPIIIEVIL 3) MAAPDRQWRRCTCRGARSTAHWKMAPGRRGRARHRSSAAYSTPHKRAQTPTCSPLRPNPPPPSALQEHHRPRRLLPHQSIRPKSSSNPSPPRRAPAGARWPASFLSVSDRRDRFPWPCLVQWLAAACCRICAAGACSPLPNRSRQWLTICPYHCSATAPVSSDGLLWP | 35 aa  32 aa  168 aa |
|  | (B) | 3 | 1) MEFELTMSSGAKMPHSIHFMEGVKLSEHGRRYMFP 2) MVSYRSIPTTFVVSSWANICMVLRPIIIEVIL 3) MAAPDRQWRRCTCRGARSTAHWKMAPGRRGRARHRSSAAYSTPHKRAQTPTCSPLRPNPPPPSALQEHHRPRRLLPHQSIRPKSSSNPSPPRRAPAGARWPASFLSVSDRRDRFPWPCLVQWLAAACCRICAAGACSPLPNRSRQWLTICPYHCSATAPVSSDGLLWP | 35 aa  32 aa  168 aa |
|  | (D) | 3 | 1) MVSYRSIPTTFVVSSWANICMVLRPIIIEVIL 2) MAAPDRQWRRCTCRGARSTAHWKMAPGRRGRARHRSSAAYSTPHKRAQTPTCSPLRPNPPPPSALQEHHRPRRLLPHQSIRPKSSSNPSPPRRAPAGARWPASFLSVSDRRDRFPWPCLVQWLAAACCRICAAGACSPLPNRSRQWLTICPYHCSATAPVSSDGLLWP 3) MSSGAKMPHSIHFMEGVKLSEHGRRRLGF | 32 aa  168 aa  29 aa |
|  | (E) | 3 | 1) MAAPDRQWRRCTCRGARSTAHWKMAPGRRGRARHRSSAAYSTPHKRAQTPTCSPLRPNPPPPSALQEHHRPRRLLPHQSIRPKSSSNPSPPRRAPAGARWPASFLSVSDRRDRFPWPCLVQWLAAACCRICAAGACSPLPNRSRQWLTICPYHCSATAPVSSDGLLWP 2) MSSGAKMPHSIHFMEGVKLSEHGRRYMFP 3) MVSYRSIPTTFVVSSWANICMVLRPIIIEVIL | 168 aa  29 aa  32 aa |

Table S3. List of primers used in experiments.

| **Primer** | **Sequence (5’ 🡪 3’)** | **Application** |
| --- | --- | --- |
| OG/8 | GCACCATGCTGGTGTAGTGGACACAAA | 5’ RACE PCR, *MIR444a* mRNA |
| OG/2 | GGTGCCACCACATGCAATAACTCAAACC | 5’ RACE PCR, *MIR444b* mRNA |
| OG/14 | GAAACGCCGACCCGGAGGAG | 5’ RACE PCR, *MIR444c* mRNA |
| OG/18 | GGCCAAAGCAAGCCATCAGATGATACG |  |
| OG650 | TTGTTGTCTCAAGCTTGCTGCCTCC |  |
| OG651 | ATAGTTCTGGCAAAGGGAGGCAGCA |  |
| OG654 | CCAAAGCAAGCCATCAGATGATACGG |  |
| OG655 | AGCGGTGGCTGAGCAATGATATGG |  |
| OG/9 | TGTTTGGGCTCTGCTTCAGTCCTCAC | 3’ RACE PCR, *MIR444a* mRNA |
| OG/15 | ATGGCTGCCCCCGATCGGCAGTG | 3’ RACE PCR, *MIR444c* mRNA |
| OG648 | AAGTGGAGGCGGCAAGCTAGAGACA |  |
| OG649 | AGGCGGCAAGCTAGAGACAGCAACT |  |
| OG653 | CCCTCCCAAATCGTAGTCGCCAAT |  |
| OG652 | GCATCTGTGCAGCTGGTGCATGTT |  |
| M13F | CACGACGTTGTAAAACGAC | Colony PCR (pGEM-T Easy) |
| M13R | GGATAACAATTTCACACAGG |  |
| APO387 | CGTGACGCTGTGTTGCTTGT | RT-qPCR, *ARF1* |
| APO388 | CCGCATTCATCGCATTAGG |  |
| OG406MIR444 | ATCAGCGCTCGCTCGCAC | RT-PCR, pri-miRNA444a |
| OG407MIR444 | AGATATTCCTTCCGTGAAGAAA |  |
| OG410MIR444 | CTCCGACTTTTCTTTCCCTCG | RT-PCR, pri-miRNA444b |
| OG411MIR444 | TCAATCACGCATCTTCTCATTTT |  |
| AC210 MIR444 | CCTCGTTCCTCTCGGTTTC | RT-PCR, pri-miRNA444c |
| AC211 MIR444 | AGGTCGTCGTCCCTTCTACA |  |
| AC2600 | GTTTGGGCTCTGCTTCAGTC | RT-qPCR, pri-miRNA444a.1 |
| AC2601 | CGAGTCAAAGAGTAGAACTGCAAA |  |
| OG1168 | AAACATAAATCTTAGGCAGCTACTCC | RT-qPCR, pri-miRNA444a.5 |
| OG1169 | CAAGAAGTTGTACTAAACTGCGACA |  |
| OG1038 | GCGGAGGACTCGAGTTTAGT | RT-qPCR, pri-miRNA444a.7 |
| OG1039 | TACTCTGAAGATACTCCCTATTTTTCC |  |
| OG1118 | AGGACTCGAGGGTCCCAAC | RT-qPCR, pri-miRNA444a.8 |
| OG1119 | CCAGGCAGCTTTGGAATAAC |  |
| OG1120 | GCGAATGAATGATCATACGAATAAG | RT-qPCR, pri-miRNA444b.1 |
| OG1121 | AGAGTGAATAGTTTAGAGTCGAATGTC |  |
| OG1058 | TGTTCCTCCAATCGCTGAAT | RT-qPCR, pri-miRNA444b.2 |
| OG1059 | AGTTTCAGGGAGGCTGGACT |  |
| OG1056 | CGAGGGTTTGAGGTAAATACTGG | RT-qPCR, pri-miRNA444b.5 |
| OG1057 | TTTAAGGTGTGGAGGGGTGAC |  |
| OG658 | TGCAGTACTTGTGGGGAAGG | RT-qPCR, pri-miRNA444c.1 |
| OG659 | AGTATGGTGATGTTCTATTAATTTTGC |  |
| OSx7 | CTCATCTGCTCCCGATGAGC | RT-qPCR, pri-miRNA444c.2 |
| OSx9 | AAAATGAAATGTAAGACCAACAGGA |  |
| OSx5 | GTCGGCGTTTCAGATGAGC | RT-qPCR, pri-miRNA444c.4 |
| OSx6 | CAGAATCCTAACCTCCTGCGC |  |
| OG692 | TGCAGTACTTGTGGGGAAG | RT-qPCR, pri-miRNA444a |
| OG693 | AGTATGGTGATGTTCTATTAATTTTG |  |
| OG660 | GGCAACAACTGCATTACTTTCA | RT-qPCR, pri-miRNA444b |
| OG661 | CGATTATGAAGACCCATAGATTTTG |  |
| OG662 | AGCAACTGCATAATTTGCAAGAA | RT-qPCR, pri-miRNA444c |
| OG664 | ATCATCGGTATGGCCCGAAA |  |
| Zpep1for | ACAGGGTCCCAACTTTGACTTTAT | RT-PCR, pri-miRNA444a, polysome fractions |
| Zpep1rev | GCTGGTGTAGTGGACACAAAAATA |  |
| 1264 | TGCTTGGTGCCACCACAT | RT-PCR, pri-miRNA444b, polysome fractions |
| 2613 | TCCGCTCGTAGATCCTTCAC |  |
| ORF168For | ATGGCTGCCCCCGATCGG | RT-PCR, pri-miRNA444c, polysome fractions |
| ORF168Rev | TCAAGGCCAAAGCAAGCCATCAGA |  |
| ADP56-R | TGTAAAACCACGGCACAGAA | RT-PCR, *ARF1* mRNA |
| ADP57-F | CTTGAAGCGTATCGAGGAC |  |
| AS104 | TCGGTCTCGTCATCTTCTCC | RT-qPCR, *MADS27* mRNA |
| AS105 | TTCGCTCGGCCATATCGATC |  |
| AS916 | TGTACCTTGCGCTGTACCTG | RT-qPCR, *NRT1.1* mRNA |
| AS917 | ACTTCTCGGTGCTGTTGGAC |  |
| Pep1HaloFor | ATCGATCGGGATGTTTGGGCTCTGCTTCAGTC | RT-PCR, PEP444a coding sequence |
| Pep1HaloRev | ATAAGAATGCGGCCGCCTAAATTGCACCATGCTGGTGTAGTG |  |
| Pep2HaloFor | ATCGATCGGGATGGAGTCCTCGTCCGCTC | RT-PCR, PEP444b coding sequence |
| Pep2HaloRev | ATAAGAATGCGGCCGCTCATGCTTGGTGCCACCA |  |
| Pep3HaloFor | ATCGATCGGGATGGCTGCCCCCGATCGG | RT-PCR, PEP444c coding sequence |
| Pep3HaloRev | ATAAGAATGCGGCCGCTCAAGGCCAAAGCAAGCCATCAGA |  |
| PS436-F | TGCAAGAAGACAACCCGGACC | Colony PCR (pH6HTN His6 HaloTaq®T7 vector) |
| PS437-R | CCCCTCAAGACCCGTTTAGAGG |  |
| orf444.1_1_ex-F | TGGCAAGACATGCCCCTTGGTAGC | sgRNA1 |
| orf444.1_1_ex-R | AAACGCTACCAAGGGGCATGTCTT |  |
| orf444.1_2_ex-F | TGGCACAAAAGTGCCAGGCAGCTT | sgRNA2 |
| orf444.1_2_ex-R | AAACAAGCTGCCTGGGACTTTTGT |  |
| orf444.2_2_ex1-F | TGGCAAACCCTCGGCGCATGCAGG | sgRNA3 |
| orf444.2_2_ex1-R | AAACCCTGCATGCGCCGAGGGTTT |  |
| orf444.2_3_ex1-F | TGGCAGCCGGGAGTGCTTGGTCTT | sgRNA4 |
| orf444.2_3_ex1-R | AAACAAGACCAAGCACTCCCGGCT |  |
| orf444.3_4-F | TGGCGTGGGGAGTAGAATATGCGG | sgRNA5 |
| orf444.3_4-R | AAACCCGCATATTCTACTCCCCAC |  |
| orf444.3_5-F | TGGCGGGGCGGTGGTGCTCTTGTA | sgRNA6 |
| orf444.3_5-R | AAACTACAAGAGCACCACCGCCCC |  |
| IK70 | GCTCACATGTTCTTTCCTGCG | Colony PCR (pIK vectors) |
| IK71 | CACCTGACGTCTAAGAAACC |  |
| Wpep2for | AAAATAAGCGGTAAATTCCAGGAT | Genotyping, *miPEP444b* |
| Wpep2rev | TATACTATTGCTCCCGCTAGGGTTC |  |
| Wpep3for | AGCACCGCCCACTGGAAAAT | Genotyping, *PEP444c* |
| Wpep3rev | ATCGATCTCTCCGATCCCGAAACC |  |
